# Imaging the columnar functional organization of human area MT+ to axis-of-motion stimuli using VASO at 7 Tesla

**DOI:** 10.1101/2022.07.29.502034

**Authors:** Alessandra Pizzuti, Laurentius (Renzo) Huber, Omer Faruk Gulban, Amaia Benitez-Andonegui, Judith Peters, Rainer Goebel

## Abstract

Cortical columns of direction-selective neurons in the motion sensitive area (MT) have been successfully established as a microscopic feature of the neocortex in animals. The same property has been investigated at mesoscale (<1 mm) in the homologous brain area (hMT+, V5) in living humans by using ultra-high field functional magnetic resonance imaging (fMRI). Despite the reproducibility of the selective response to axis-of-motion stimuli, clear quantitative evidence for the columnar organization of hMT+ is still lacking. Using cerebral blood volume (CBV)-sensitive fMRI at 7 Tesla with submillimeter resolution and high spatial specificity to microvasculature, we investigate the columnar functional organization of hMT+ in 5 participants perceiving axis-of-motion stimuli for both blood oxygenation level dependent (BOLD) and vascular space occupancy (VASO) contrast mechanisms provided by the used Slab-Selective Slice Inversion (SS-SI)-VASO sequence. With the development of a new searchlight algorithm for column detection, we provide the first quantitative columnarity map that characterizes the entire 3D hMT+ volume. Using voxel-wise measures of sensitivity and specificity, we demonstrate the advantage of using CBV-sensitive fMRI to detect mesoscopic cortical features by revealing higher specificity of axis-of-motion cortical columns for VASO as compared to BOLD contrast. These voxel-wise metrics also provide further insights on how to mitigate the highly debated draining veins effect. We conclude that using CBV-VASO fMRI together with voxel-wise measurements of sensitivity, specificity and columnarity offers a promising avenue to quantify the mesoscopic organization of hMT+ with respect to axis-of-motion stimuli. Furthermore, our approach and methodological developments are generalizable and applicable to other human brain areas where similar mesoscopic research questions are addressed.

## 1. Introduction

An important task of the visual system is to recognize moving objects in a dynamic environment. The macaque middle temporal (MT) area and its homologue in humans (hMT+, V5) is a higher order cortical visual area sensitive to the direction of motion (Watson et al., 1993; Born and Bradley, 2005; Zeki, 1974; Maunsell and Van Essen, 1983; Rees et al., 2000; Zimmermann et al., 2011). It also plays an important role in constructive motion perception as revealed by illusory (e.g. apparent) motion stimuli (Goebel et al., 1998; Muckli et al., 2002; Born and Bradley, 2005; Schneider et al., 2019). Invasive electrophysiological recordings (Albright et al., 1984; Diogo et al., 2003) and optical imaging (Malonek et al., 1994) results suggested that macaque extra-striate area MT is functionally organized in cortical columns, similarly to V1 (Hubel and Wiesel, 1965). In particular, Albright et al. (1984) demonstrated in macaque MT the occurrence of a systematic relationship between the penetration angle of the electrode and the rate of change of preferred direction of motion, indicating that cells with a similar preference are arranged in vertically oriented columns (Mountcastle, 1997; Buxhoeveden and Casanova, 2002). Albright et al. also discovered that opposing directions of motion are located in neighboring columns forming larger axis-of-motion columns.

The advent of magnetic resonance imaging opened the avenue to bridge the gap to animal literature, by non-invasively investigating the functional organization of the human brain. Especially the advent of ultra high field (UHF) functional magnetic resonance imaging (fMRI) combined with recent advances in MR hardware and pulse sequences offers sub-millimeter spatial resolution (Uğurbil et al., 2003; Shmuel et al., 2007; Uludağ et al., 2009; Koopmans and Yacoub, 2019) providing a unique way to unveil mesoscopic properties underlying cognitive functions at the level of cortical layers and cortical columns (Rakic, 2008; Petro and Muckli, 2017; Uğurbil, 2016; Dumoulin et al., 2018; De Martino et al., 2018; Viessmann and Polimeni, 2021; Uğurbil, 2021; Huber et al., 2020; Cho et al., 2022). Mesoscopic 7T fMRI (Zimmermann et al., 2011; De Martino et al., 2013) has already been used to investigate the columnar organization of human MT using 3D-GRASE (Oshio and Feinberg, 1992; Feinberg et al., 1995; Kemper et al., 2015) and BOLD contrast. Zimmermann et al. (2011) provided the first direct demonstration of axis-of-motion selectivity and tuning characteristics in the human brain together with first insights into the spatial organization of hMT. However, their columnar results can only be seen as an early approximation, since their limited number of samples across cortical depth could obscure the full arrangement of the vertical columnar property. To validate and enhance previous findings, we investigate the functional organization of hMT+ in humans perceiving axis-of-motion stimuli using three improvements: (I) we use cerebral blood volume (CBV)-sensitive fMRI in addition to conventional BOLD fMRI, and we developed novel methods to (II) quantify local sensitivity and specificity, and (III) to detect, characterize and visualize cortical columns.

### CBV-sensitive fMRI

We use the SS-SI VASO (Huber et al., 2014) pulse sequence to get high-resolution CBV-sensitive fMRI. The VASO sequence is more suitable for imaging mesoscopic features compared to standard gradient echo BOLD echo planar imaging (GE-BOLD EPI), since it provides higher spatial specificity to the neuronal activation cite (Huber et al., 2019). Differently from the BOLD contrast, VASO is more sensitive to the microvasculature and it is less compromised by both oxygenation related vascular changes as well as large draining effects from vessels penetrating the cortex orthogonally or from pial veins lying on top of the gray matter surface (Duvernoy et al., 1981; Weber et al., 2008; Lauwers et al., 2008). However, its usage at submillimeter resolution is still challenging due to its inherently low signal-to-noise ratio (SNR) (Huber et al., 2019).

### Local measure of sensitivity and specificity

For more precise quantification of differences between BOLD and VASO fMRI responses, we develop a voxel-wise measure of sensitivity and specificity as an extended version of the conventional global measures (Beckett et al., 2020; Huber et al., 2017). These metrics were also used to compare results from BOLD and VASO contrast mechanisms at the global level (Huber et al., 2017, 2020; Oliveira et al., 2022). Global metrics summarize the behavior of an entire region of interest, whereas our local metrics provide a functional characterization for each voxel inside a region of interest, resulting in more complete information for addressing mesoscopic questions at submillimeter resolution. More specifically, our new local metrics allow to evaluate the tuning property encoded in each voxel: if a voxel is strongly tuned to a specific condition its specificity will be high and its sensitivity will be inherited by the contrast mechanism, whereas if a voxel is poorly tuned, as in the case of a voxel sampling a big vessel, the specificity will be very low while the sensitivity will be very high.

### Novel method to characterize cortical columns

To overcome depth-sampling limitations (Zimmermann et al., 2011), we develop a new searchlight algorithm for functional column detection that inspects the whole cortical depth of a specific volume of interest and provides a quantitative columnarity map that is useful to detect highly-columnar functional patches inside the volume. The algorithm provides a new generalized framework for investigating the columnar functional organization for any cortical gray matter ribbon and it can potentially improve the replicability and comparability of columnar results across studies using a variety of columnar quantification methods (Yacoub et al., 2008; Zimmermann et al., 2011; De Martino et al., 2013, 2015; Schneider et al., 2019).

In this study we acquire ultra-high field (7 Tesla) fMRI at submillimeter resolution (0.8 mm isotropic) to investigate the axis-of-motion cortical columns in hMT+ (V5). By using our new algorithm for column detection, we provide a quantitative columnarity map that fully characterizes the functional organization of hMT+ with respect to axis-of-motion preference. We demonstrate the advantage of using CBV-sensitive fMRI to detect mesoscopic cortical features by showing the higher specificity of axis-of-motion cortical columns detected by VASO compared to BOLD contrast.

## 2. Materials and Methods

### 2.1. Experimental design

#### Participants

Five healthy participants (4 males and 1 female, 28-34 years old, 10 hMT+ regions) with normal or corrected-to-normal vision were recruited for the study. Participants received a monetary reward. All participants had been in an MRI scanner at least once before and were trained and experienced at maintaining fixation for long periods of time. Informed consent was obtained from each participant before conducting the experiment. The study was approved by the ethics review committee of the Faculty of Psychology and Neuroscience (ERCPN) of Maastricht University and experimental procedures followed the principles expressed in the Declaration of Helsinki.

#### Stimulus presentation

The scripts used for the stimulus presentation were developed based on Schneider et al. (2019) and presented using the open source application PsychoPy3 (v2020.2.4). Scripts are available on https://github.com/27-apizzuti/AOM-VASO_project. A frosted screen (distance from eye to screen: 99 cm; image width: 28 cm; image height: 17.5 cm) at the rear of the magnet was used to project the visual stimuli (using Panasonic projector 28 PT-EZ570; Newark, NJ, USA; resolution 1920 × 1200; nominal refresh rate: 60 Hz) that participants could watch through a tilted mirror attached to the head coil. We used 50% gray background (at 435 cd/m^2^ luminance) with white dots (at 1310 cd/m^2^) for the motion stimulation (black color is measured at 2.20 cd/m^2^). An MR compatible button box was used to register participants’ responses for the attention task during the entire functional scanning procedure.

#### Stimulus description

Each participant underwent one 2 hour scanning session. In the scanning session, we collected 1 run to functionally locate hMT+, 3-4 runs (according to the available scanning time) to map four axes of motion conditions, and an MP2RAGE scan to obtain a high-resolution structural image (only if not already available from a previous scanning session). For the hMT+ functional localizer, a standard block design paradigm was used presenting moving dots in alternation with static dots in a circular aperture (Tootell et al., 1995) (run duration: 9 min 10 s). We did not separate MST from hMT (Huk et al., 2002; Kolster et al., 2010; Zimmermann et al., 2011) since we aimed to cover the whole human motion complex (hMT+). Dots traveled inwards and outwards from the center of the aperture for 10 seconds (speed = 8 degree of visual angle per second, dot size = 0.2 degree of visual angle, number of dots = 200, black dots on gray background), were followed by a stationary dots display presented for the same amount of time. A total of 27 repetitions of task-rest blocks were collected. For the axis-of-motion mapping runs, we presented dots in a circular aperture moving coherently along one of four axes (0° ↔ 180°, 45° ↔ 225°, 90° ↔ 270°, 135° ↔ 315°, in both directions of motion) (Zimmermann et al., 2011) alternated with static flickering dots (run duration: 15 min). Moving dot patterns (speed = 8 degrees of visual angle per second, dot size = 0.2 degree of visual angle, number of dots = 250) were presented for 24 seconds followed by a variable inter-trial interval (ITI) of 24–29 seconds to reduce functional signal carry over effects. Each motion axis block was repeated four times per run. In all conditions, a black disk (target/fixation dot) surrounded by an annulus was presented in the center of the aperture. Participants were instructed to fixate the black disk and respond through the button box every time the annulus changed color (attention task). A demo of our axis-of-motion stimulus is available here: https://github.com/27-apizzuti/AOM-VASO_project/tree/main/stimulus_scripts/demos.

### 2.2. MRI acquisition

Data acquisition was performed on a whole-body “classical” MAGNETOM 7T (Siemens Healthineers, Erlangen, Germany) at Scannexus B.V. (Maastricht, The Netherlands) using a 32-channel RX head-coil (Nova Medical, Wilmington, MA, USA). A second and a third order B0 shimming procedure was used to improve the homogeneity of the main magnetic field B0 in the shim volume containing the region of interest. The localizer experiment was conducted using a GE EPI sequence with BOLD contrast (Moeller et al., 2010) (echo time (TE)=15 ms, nominal flip angle (FA)=55°, echo repetition time (TR)=1000 ms, multi band factor (MB)=3, 57 slices) with a whole brain field of view and a (98×98) matrix at 2 mm isotropic nominal resolution. Before the acquisition of the run, we collected 5 volumes for distortion correction with the settings specified above but opposite phase encoding (posterior-anterior).

For the axis-of-motion mapping experiment, we used a Vascular Space Occupancy (VASO) sequence (Lu et al., 2003) optimized for 7T (Hua et al., 2013). Specifically, we used the Slab-Selective Slice Inversion (SS-SI VASO) approach (Huber et al., 2014) with a 3D EPI readout (Poser et al., 2010) at nominal isotropic voxel resolution of 0.8 mm. Previous work has shown that the 3D readout is beneficial for sub-millimeter applications (Huber et al., 2018). The in-plane field of view was 129×172 mm (162×216 matrix) for a total of 26 acquired slices. The imaging parameters were: TE=25 ms, ‘temporal resolution’ of pairs of images=4840 ms, variable flip angle scheme FA=26+°, in plane partial Fourier factor 6/8 with POCS reconstruction of 8 iterations, inversion time (TI)=1530 ms for the blood nulling point and FLASH-GRAPPA=3 (Talagala et al., 2016). Variable flip angles were used to minimize T_1_-related blurring along the slice direction (Huber et al., 2018). The sequence was implemented using the vendor provided IDEA environment (VB17A-UHF) and is available to download via C2P on the SIEMENS App-Store in Teamplay. The placement of the small functional slab was guided by an online analysis of the hMT+ localizer data (general linear modeling by Siemens), to ensure a bilateral coverage of area hMT+ for every participant.

The anatomical images were acquired with an MP2RAGE (magnetization prepared two rapid gradient echoes) (Marques et al., 2009) at 0.7 mm isotropic resolution (TR/TE=6000 ms/2.39 ms, TI=800 ms/2750 ms, FA=4°/5°, GRAPPA=3). MP2RAGE sequence parameters are optimized to overcome the large spatial inhomogeneity in the transmit B1 field by generating and combining in a novel fashion two different images at two different inversion times (TI1, TI2) to create T_1_-weighted MP2RAGE uniform (UNI) images (Marques et al., 2009). Physiological traces of respiration and heartbeat were recorded but not used for the analysis, since we did not observe any improvements by conducting `RETROICOR` physiological noise correction during the piloting stage (Glover et al., 2000). Indeed, sub-millimeter data are expected to be in the thermal noise-dominated regime (Triantafyllou et al., 2005).

### 2.3. Structural data analysis

#### 2.3.1. Preprocessing and registration

T_1_-weighted UNI images with high contrast-to-noise ratio from MP2RAGE were used to guide the determination of the anatomical location of hMT+ and to derive layers in the cortical ribbon of interest. T_1_-w UNI images were skull-stripped using a brain mask obtained by inputting the MP2RAGE INV2 (TI2) images to FSL BET (v.6.0.5) (Smith et al., 2004). When needed, we semi-automatically corrected the results and used Segmentator (v.1.6.0) (Gulban et al., 2018) to remove dura mater contribution. Then, we used the anatomical bias correction algorithm from SPM12 (Wellcome Trust Center for Neuroimaging, London, UK) to further reduce inhomogeneities in the T_1_-w UNI images (Friston et al., 2007; Ashburner and Friston, 2005). In order to preserve the high-resolution functional data from issues of registration and interpolation while computing depth-dependent analysis (Huber et al., 2017; Guidi et al., 2020), we aligned the T_1_-w UNI images to the high-resolution functional VASO data, according to https://layerfmri.com/2019/02/11/high-quality-registration/. The target functional slab “T_1_-w EPI” was derived from the original SS-SI VASO time series by computing the inverse of signal variability that provides a good contrast between gray matter (GM) and white matter (WM) using AFNI (v.20.3.01) (Cox, 1996; Cox and Hyde, 1997) (function: `-cvar`). In ITK-SNAP (v.3.8) (Yushkevich et al., 2006) we manually aligned the T_1_-w UNI whole brain images to the T_1_-w EPI slab and then ran ITK-SNAP’s automatic co-registration tool. Then, we used the obtained transformation matrix as input for running ANTS’s “Syn” registration algorithm (v.20.3.01) (Avants et al., 2009, 2011; Madge, 2020). Finally, the T_1_-w UNI images were resampled to the T_1_-w EPI space using ANTS b-spline interpolation (`antsApplyTransforms -BSpline`).

#### 2.3.2. Segmentation

Accurate and precise tissue segmentation is a crucial step to investigate cortical layers and columns. Since conventional segmentation packages yielded unsatisfactory results for our restricted field of view, a semi-automated segmentation approach with manual intervention was used here. We used `-3dresample` command from AFNI (v.20.3.01) to upsample (with cubic interpolation) the processed T_1_-w UNI images with an upscaling factor of 4 (nominal resolution = 0.2 mm isotropic). This step allowed a smoothed calculation of cortical features (e.g. curvature or thickness). Then, we confined the segmentation process to a “scoop of interest” in both hemispheres: in ITK-SNAP we centered a spherical mask around the anatomically expected area hMT+. We ran FSL FAST to obtain a first definition of the cerebro-spinal fluid/ gray matter (CSF/GM) and the gray matter/white matter (GM/WM) tissue borderlines. Tissue labels were carefully quality controlled and manually edited when necessary (by A.P.), and later revised independently by another expert (O.F.G.). In combination, we also used morphological operations (dilation and erosion) (Virtanen et al., 2020) to further improve the segmentation output. These operations, when applied in combination with the same parameters, remove mislabeled isolated voxels and smooth boundaries between tissues.

#### 2.3.3. Cortical depths

Once the segmentation was completed, we used `LN2_LAYERS` program from LayNii (v2.2.1) (Huber et al., 2021) to compute equi-volume cortical depths (Bok, 1959; Waehnert et al., 2014), cortical thickness and curvature for each gray matter voxel.

#### 2.3.4. Flattening

We used `LN2_LAYERS, LN2_MULTILATERATE`, and LN2_PATCH_FLATTEN programs within LayNii to flatten our cortical patches (Gulban et al., 2022). We first establish the center of gravity of the gray matter activated ROIs (left and right hMT+). Then a disk of a predefined radius is grown geodesically and a local 2D coordinate system (U and V coordinates) was imposed on it (using `LN2_MULTILATERATE` program). Together with the “metric” file from `LN2_LAYERS` (D coordinate), the end result of this procedure is a full continuous mapping between flat cortex space (UVD) and the original folded cortex space (XYZ). This mapping allows us to flatten 3D chunks of cortical data (in NIfTI format) to explore mesoscopic cortical structures as it was done in previous studies (Zimmermann et al., 2011; De Martino et al., 2013; Schneider et al., 2019).

### 2.4. Functional data analysis: Localizer experiment

Functional localizer data were pre-processed in BrainVoyager v.22.1 (Goebel et al., 2006, 2012) as follows: slice scan time correction, motion-correction, linear trend removal and high-pass filtering (6 cycles). We corrected for EPI geometric distortion using the image registration method based on the opposite phase encoding direction EPI data as implemented in COPE BrainVoyager plugin (Breman et al., 2020). The same aligning procedure explained for structural images was also used to align functional localizer data to high-resolution VASO data. The registration parameters (ITK-SNAP, ANTS “Syn”) were estimated matching the computed temporal mean image from localizer data (Smith et al., 2004) with T_1_-w EPI and then applied to the time series using ANTS B-spline interpolation. Then, in order to functionally define our ROI (bilateral hMT+), we calculated a voxel-wise general linear model (GLM) for each participant in BrainVoyager. The GLM was corrected for temporal auto-correlation (AR2). The model contained a single predictor for the stimulus condition “moving dots” convolved with a standard hemodynamic response function. Voxels that showed a significant response to “moving vs static dots” contrast (using a threshold (q) corrected for multiple comparisons using false discovery rate; q(FDR) < .05) were selected. Finally, we defined a bilateral hMT+ ROI, by intersecting these voxels with two spheres of 16 mm in radius (one for each hemisphere) placed inside the expected anatomical location (Zimmermann et al., 2011; Schneider et al., 2019).

### 2.5. Functional data analysis: Axes of motion experiment

#### 2.5.1. Preprocessing and functional maps

Axis-of-motion functional data were analyzed following the optimized preprocessing pipeline for SS-SI VASO sequence https://layerfmri.com/2019/03/22/analysispipeline/. For each run, the first four time points (non-steady state images) were overwritten with steady-state images. Then, we separated odd and even time points from raw data, corresponding to MR signals with and without blood nulling and we separately performed motion correction using SPM12 (Friston et al., 2007). A fourth-order spline was used for resampling to minimize blurring. For every participant, motion parameters between the two contrasts were very similar, as expected, and never higher than the nominal voxel resolution. The original time series length was restored for both blood-nulled and BOLD images using 7^th^ order polynomial interpolation method before multiple runs were averaged to minimize noise amplification in the next processing steps. Dynamic division of blood-nulled and BOLD volumes was performed to generate VASO images with reduced BOLD contrast contamination (`LN_BOCO` program within LayNii). BOLD correction is valid under the assumption that T_2_* contrast is the same in images with both contrasts, because they are acquired concomitantly (Huber et al., 2014). For each participant, we fit a voxel-wise general linear model restricted to the bilateral hMT+ ROI (previously defined) on both the BOLD and VASO time series in BrainVoyager. The model contained five predictors, one for each axis-of-motion condition and one for the flickering condition. Each predictor was then convolved with a standard hemodynamic response function. The GLM was corrected for temporal auto-correlation (AR2). Statistical t-maps were then computed. While generating the t-map by contrasting one axis-of-motion stimulus condition (e.g. horizontal) versus flickering dots baseline condition, the two contrasting conditions were balanced in terms of number of time points considered in the computation, since in the experimental paradigm the latter was repeated four times more than the former one. In addition to the t-maps, we also computed voxel-wise percent signal change (Huber et al., 2017; Beckett et al., 2020) between mean signal during task (e.g. moving dots along one axis-of-motion) and baseline (flickering dots) with the same balancing rationale used for computing t-maps. This method has been shown to provide results that are easier to interpret than methods using inferential statistics, which can be affected by laminar differences in noise and hemodynamic response function shape (Huber et al., 2017). We used percent signal change to computed layer profiles (results are presented in **Supplementary Figure 2 and 3**).

#### 2.5.2. Preference maps and voxel selection

We created a BOLD and VASO “preference map” (un-thresholded) by assigning to each voxel the preferred axis-of-motion, based on the predictor showing the highest GLM fit (among the four axes of motion), representing the highest stimulus-induced normalized fMRI response (t-value) (Zimmermann et al., 2011; De Martino et al., 2013; Schneider et al., 2019).

To take into account differences in sensitivity between the two contrast mechanisms, we evaluate the hMT+’s tuning to axes of motion stimuli following three statistical ways and compare the obtained results.

We thresholded the preference maps by considering three sets of voxels (see Table1): 1) voxels that exhibited a significant t-value response (using a threshold (q) corrected for multiple comparisons using false discovery rate; q(FDR)<.05) when contrasting “all axes of motion moving dots vs flickering dots”, later called ‘FDR’. 2) Voxels that survive a cross-validation test, later called ‘CV’. 3) Voxels that survive a cross-validation test and show positive t-value response to each axis-of-motion condition, later called ‘CV+’. A cluster-size thresholding (threshold=4) was applied separately to each set of voxels (using https://gist.github.com/ofgulban/27c4491592126dce37e97c578cbf307b).

**Table 1:** Number of BOLD and VASO voxels considered for each voxel selection method (FDR, CV, CV+).

| Left-hMT+ | BOLD |  |  | VASO |  |  |
| --- | --- | --- | --- | --- | --- | --- |
|  | FDR | CV | CV+ | FDR | CV | CV+ |
| sub-01 | 1168 | 444 | 404 | 88 | 302 | 157 |
| sub-02 | 1245 | 388 | 370 | 66 | 203 | 82 |
| sub-03 | 1006 | 394 | 383 | 48 | 258 | 88 |
| sub-04 | 549 | 191 | 184 | 9 | 141 | 55 |
| sub-05 | 756 | 286 | 261 | 149 | 256 | 123 |

| Right-hMT+ | BOLD |  |  | VASO |  |  |
| --- | --- | --- | --- | --- | --- | --- |
|  | FDR | CV | CV+ | FDR | CV | CV+ |
| sub-01 | 976 | 384 | 361 | 25 | 157 | 54 |
| sub-02 | 1159 | 410 | 391 | 9 | 226 | 75 |
| sub-03 | 1082 | 439 | 430 | 40 | 262 | 107 |
| sub-04 | 419 | 133 | 127 | 4 | 74 | 25 |
| sub-05 | 1152 | 497 | 476 | 28 | 234 | 85 |

The number of ‘FDR’ voxels differs significantly between the two contrast mechanisms, due to expected differences in sensitivity. As an alternative approach to the FDR method, we designed a leave-one-run out cross-validation method (Zimmermann et al., 2011; Emmerling, 2016) that might be less dependent on the lower sensitivity of VASO. The cross-validation was separately applied to both BOLD and VASO preference maps restricted to suprathreshold BOLD ‘FDR’ voxels. Iteratively, we divided the available number of runs (learning set) into training set (average of all runs excluding the test run) and test set (left out run) and for each set we labeled voxels according to their preferred axis, by fitting a GLM as explained above. Within each fold of the cross-validation process, only voxels showing the same axis preference between training and test set were kept for the next step. Finally, we cross-validated our BOLD and VASO preference maps by removing all voxels whose labeled preferred axis did not match with the cross-validated predicted label.

Our third method tackles the question about the neural meaning of having a negative response to a specific condition that was not taken into account for FDR and CV voxels. A negative response to a specific axis-of-motion could, in principle, indicate a neural suppression, as shown here Albright (1984). However, positive and negative t-value fluctuation around zero can also occur as noise contribution when dealing with low contrast-to-noise-ratio (CNR) data as it is the case for high-resolution fMRI. In order to exclude the latter possibility, we introduced the ‘CV+’ set of voxels. In this case, we narrowed down the set of CV-voxels by only keeping those voxels that showed a positive response (t-value) for all conditions, when separately contrasting each axis-of-motion with the ‘flickering condition’.

#### 2.5.3. Tuning Curves

Finally, for each set of voxels, tuning curves are computed for each axis-of-motion, by averaging t-values responses of all voxels sharing the same preferred axis-of-motion and evaluating them in the preferred and not preferred conditions. As previously done by (De Martino et al., 2013), a global tuning selectivity index was also computed for each tuning curve, by calculating the ratio of the response for the preferred axis-of-motion divided by the average response towards all other axes of motion.

#### 2.5.4. Voxel-wise sensitivity and specificity calculation

To investigate the signal behavior and differences between BOLD and VASO contrast, we developed new voxel-wise measures of sensitivity and specificity. By coding the voxel response to each axis-of-motion with a t-value, a four entries t-value vector was assigned to each voxel. Sensitivity was computed by calculating the Euclidean norm of the t-values vector 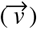 (Eq.1):

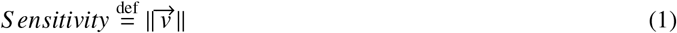

Specificity was computed according to Eq.2:

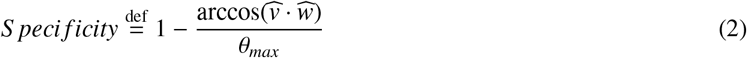

First, the t-values vector is transformed into a ‘unit vector’ by dividing each component with the norm of the vector (from v to 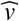). Then, the entries of the vector were arranged in ascending order and the dot product between 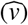 and a reference winning vector was computed [0 0 0 1] 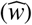. Since the ordering operation constrains the angle computation in the range 0-60° (in 4 dimensions), we scaled the computed angle by the maximum angle (*θ_max_*). Finally, we defined “specificity” as the additive inverse of the computed scaled angle.

In **Figure 1** we show with two numerical examples how to compute step-by-step our voxel-wise sensitivity and specificity measures. Note that *θ_max_* for the 2D case is equal to 45°. The sensitivity measure quantifies how strongly a voxel responds to any axis-of-motion condition. The specificity measure quantifies how well a voxel is tuned toward a specific axis-of-motion. We developed our specificity index (inspired by the n-dimensional compositional vector angle analysis from Gulban (2018)) which serves a similar purpose to the orientation selectivity index (OSI) commonly used in orientation tuning studies (Swindale, 1998; Yacoub et al., 2008; Cho et al., 2022). The OSI is used when orientation tuning is investigated with a high orientation sampling (e.g. minimally every 45 degrees) in order to detect smooth changes in orientations, to fit a circular distribution and evaluate its dispersion from the preferred orientation. However, the OSI is not suitable for axis-of-motion paradigms since they do not differentiate functional responses to opposite motion orientation and, as consequence, do not provide a continuous sampling of the tuning response. In contrast, our specificity measure does not require a fitting and therefore may be considered as an alternative way to quantify voxel selectivity even in cases when categorical (and not continuous) stimulations are evaluated. For the specificity index calculation, negative t-value entries are a critical point. We assume that negative t-values represent, in our data, an absence of neural response to a specific condition, therefore, when occurring in FDR and CV set of voxels, we zero them. The CV+ set of voxels does not suffer from this limitation and functions as control.

**Figure 1:**
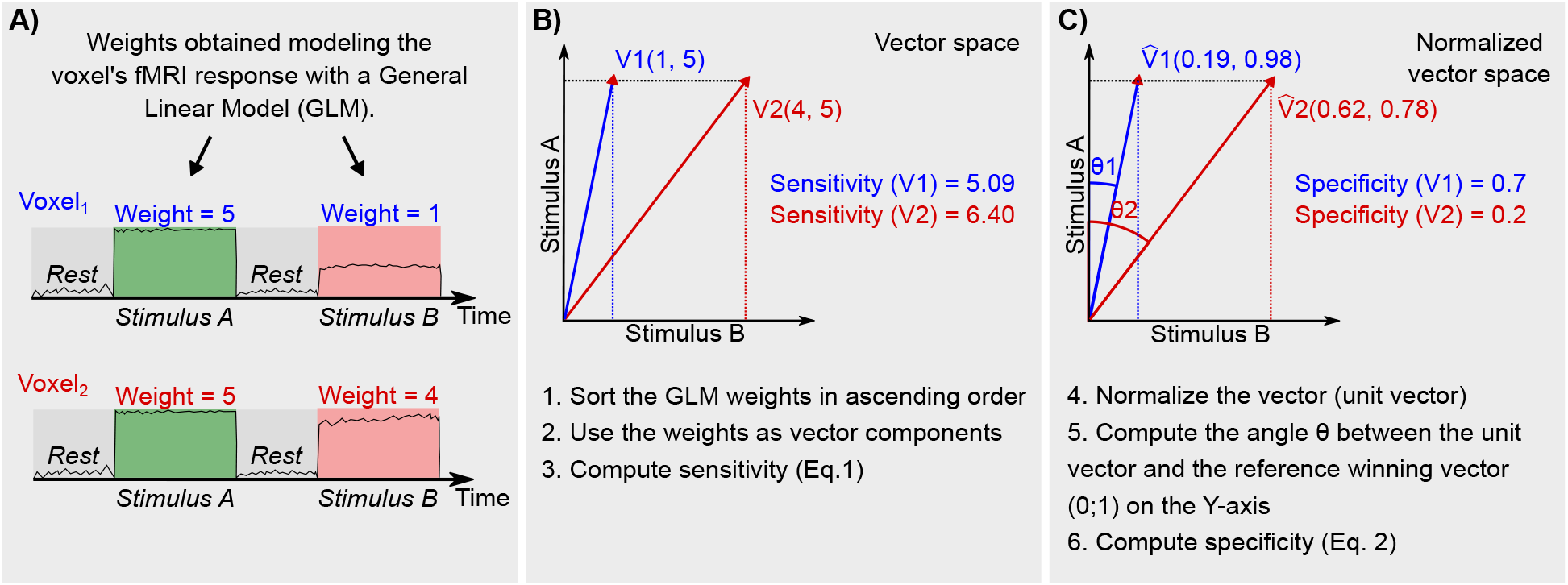
Numerical example of how to compute voxel-wise sensitivity and specificity for a 2 dimensional case. In A) the fMRI response of a voxel is modeled using a General Linear Model and the response modulation or “weight” (e.g. beta, t-value, percent signal change) during each stimulus is computed. In B) and C) we show step-by-step how to use the weights to compute our voxel-wise sensitivity and specificity.

The inset at the upper right corner of **Figure 4A** visualizes the relationships between these two variables: voxel characteristics strongly differ according to which region of the space they belong to. For instance, high-tuned voxels are represented with high specificity, whereas un-specific vessel-dominated voxels are represented by high sensitivity and low specificity.

To test if it is possible to derive information about the vasculature from our fMRI data, we combined sensitivity and specificity information with time-averaged T_2_*-weighted EPI intensity from BOLD time series, since it was demonstrated to be a robust marker of vascular effects (Kay et al., 2019). Global measures of sensitivity and specificity were also implemented to compare our results to the literature (Beckett et al., 2020; Huber et al., 2017).

#### 2.5.5. Voxel-wise columnarity index calculation

Cortical column detection and quantification using fMRI is a challenging task (Yacoub et al., 2008; Zimmermann et al., 2011; De Martino et al., 2015; Schneider et al., 2019). Inspired by Blazejewska et al. (2019), we implemented a searchlight algorithm to seek cylindrical columnar structures following the local coordinates of the cortical gray matter. This algorithm is implemented as a part of `LN2_UVD_FILTER` program (accessible via “-columns” option) in LayNii v2.2.1 and consists of the following steps:

1. Our primary inputs are the local coordinates of the cortical gray matter. These local coordinates consists of: (i) a voxel-wise parametrization of the cortical surface that contains the orthogonal U and V coordinates (computed by `LN2_MULTILATERATE` program), and (ii) voxel-wise equi-volume cortical depths that contains the D coordinates (“metric” file output as computed by `LN2_LAYERS` program) (see **Figure 2A**).
2. Our secondary input is a scalar map. This scalar map consists of the BOLD or VASO preference map (see **Figure 2B**).
3. Our tertiary input is a binary map which we refer to as “domain”. This binary map consists of the initial set of tuned voxels (the “FDR BOLD” voxels were used as mask for both BOLD and VASO contrast, see definition in the method section “Preference maps and voxel selection”). Note that we extended these activated voxels to cover the rest of the cortical thickness (using `LN2_UVD_FILTER` with “-max” option). For instance, if only a single middle gray matter voxel is available at a location, we include the voxels above and below until it covers the local cortical thickness.
4. The algorithm starts by evaluating a cylindrical 3D window centered at every voxel within the domain. This evaluation is done by (i) computing Euclidean distances using the UVD coordinates of the center voxel to all the other voxels within the domain, and (ii) by detecting the voxels that fall within user-determined radius and height of the cylinder. Note that we set the cylinder radius to 0.6 mm and the height to cover the full cortical depth (independent of a voxel being at the deep, middle, or superficial part of the cortex, see number I, II, III, IV in **Figure 2C**). We chose the diameter of the cylinder to be slightly higher than the nominal resolution (0.8 mm) in order to take into account all the possible orientations that a voxel can assume to sample the brain geometry (our diameter 1.2 mm is in the range: 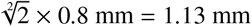 and 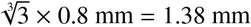).
5. This step determines a set of voxels for each window. Note that the number of windows is equal to the number of voxels within the domain input. However, the total number of voxels for each window can change as a function of local cortical thickness and local curvature. Note that this behavior is natural and expected.
6. For the set of voxels within each window, we compute the modal (most frequently occurring) value within the BOLD or VASO preference maps.
7. Finally, we compute the “columnarity index” by dividing the number of voxels of the modal value with the total number of voxels evaluated in the cylindrical window (see **Figure 2D**). Note that the columnarity index ranges between 0 (no columnarity) to 1 (ideal or pure columnarity).

**Figure 2:**
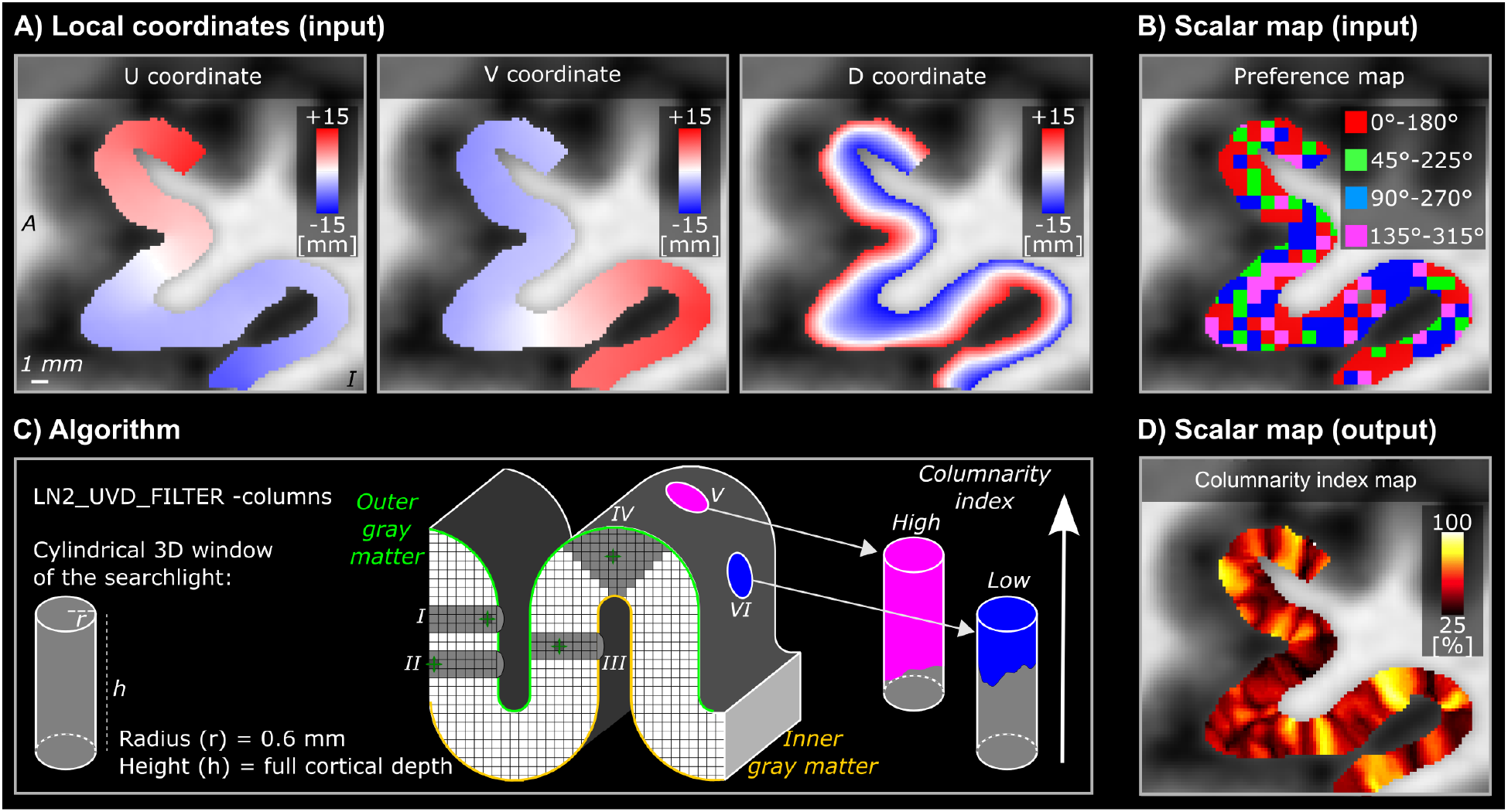
Overview of the searchlight algorithm for functional cortical column detection. A-B) Input examples of local coordinates (U, V, D) and BOLD preference map of left-hMT+ for one example participant (sub-01). C) Conceptualization of the algorithm on a toy model of a cortical ribbon. A searchlight with a cylindrical 3D window is evaluated at each voxel position. On the toy model, we show the searchlight (dark gray voxels with cardinal axes) and its relative 3D window (light gray). For every position of the searchlight, the window adapts to the geometry of the ribbon and covers the entire cortical depth (see examples (I) voxel close to the outer gray matter, (II) voxel close to the inner gray matter, (III) voxel close to the middle gray matter on a wall, (IV) voxel close to the middle gray matter on a gyrus). Pink cylinder is an example of a high columnarity index (V), whereas the blue cylinder is an example of a low columnarity index (VI). D) Output example of the columnarity index map generated for the data (B) of the same participant.

Upon completion, our algorithm yields a map where each voxel contains a unique columnarity index. This procedure allows us to reveal cortical columns for any range of the columnarity index (from 0 to 1, with 0.5 step size) by accordingly thresholding the columnarity map and applying it as a binary mask to the input preference map.

We summarize the columnar behavior of hMT+ for both BOLD and VASO contrast by providing the following quantitative measurements as a function of the columnarity index: voxel-wise sensitivity and specificity (see **Figure 8A,B (i-ii)**) and percentage of columnar volume with respect to the volume of the domain (see **Figure 8A,B (iii)**). Finally, we evaluate the spatial similarity between BOLD and VASO columnarity maps by computing a spatial consistency score: for each value of the columnarity index we quantify the percentage of voxels that appear in the same spatial location and with the same axis-of-motion preference in both contrasts with respect to the total amount of columnar voxels (see **Figure 8A,B (iv)**).

#### 2.5.6. Control analysis: benchmarking columnarity index calculation

In order to interpret the columnar results from the empirical BOLD and VASO functional data in terms of vicinity to a ‘pure noise’ or to an ‘ideal columnar’ scenario, we ran our columnarity index computation on two synthetic scalar maps (‘noise’ preference map and ‘ideal’ preference map) and compared empirical vs benchmark results. This procedure was done for each empirical hMT+ separately (i.e., 10 hMT+ regions from the left and right hemisphere of five participants). The ‘random’ preference map was generated at original voxel resolution (0.8 iso mm) by assigning to each voxel a preferred axis-of-motion based on a random distribution and then upsampled, as done for the empirical data. The ‘ideal’ preference map that would lead to ‘ideal’ functional columns was created by using the LN2_HEXBIN program from Laynii. This program uses the U,V coordinates of a specific brain volume, to generate hexagonal bins with a desired radius spanning for the whole cortical depth. The hexagonal bins are considered as an approximation of ‘ideal’ geometrically-defined cortical columns. We chose 1 mm diameter to simulate columns that follow our hypothesis of axis-of-motion columns that can be unveiled by our current resolution. We compared columnarity index distributions from our empirical and benchmark data (results are presented in **Figure 9** and in **Supplementary Figure 4**).

## 3. Results

### 3.1. Axes-of-motion tuning curves confirm direction-selectivity of hMT+

Participants performed well (97% accuracy) on the change detection task, indicating proper fixation throughout the experimental runs. The tuning property of hMT+ with respect to axis-of-motion stimuli was evaluated separately for both BOLD and VASO in three sets of voxels (FDR, CV, CV+). In **Table 1**, we report for each participant and hemisphere the number of voxels belonging to each set. We found bilateral axes-of-motion specific tuning curves for all three sets of voxels (FDR, CV, CV+) for both the BOLD and VASO contrast. As can be seen in **Figure 3**, a characteristic peak at the preferred axis-of-motion is present in each group of voxels, showing a clear preference for a single axis-of-motion. Here we show, for the first time, that VASO is also sufficiently sensitive to map differential axes of motion responses in hMT+. Moreover, our BOLD results corroborate and extend previous fMRI BOLD findings (Zimmermann et al., 2011; De Martino et al., 2013; Schneider et al., 2019) showing robust responses of hMT+ to directions of motion.

**Figure 3:**
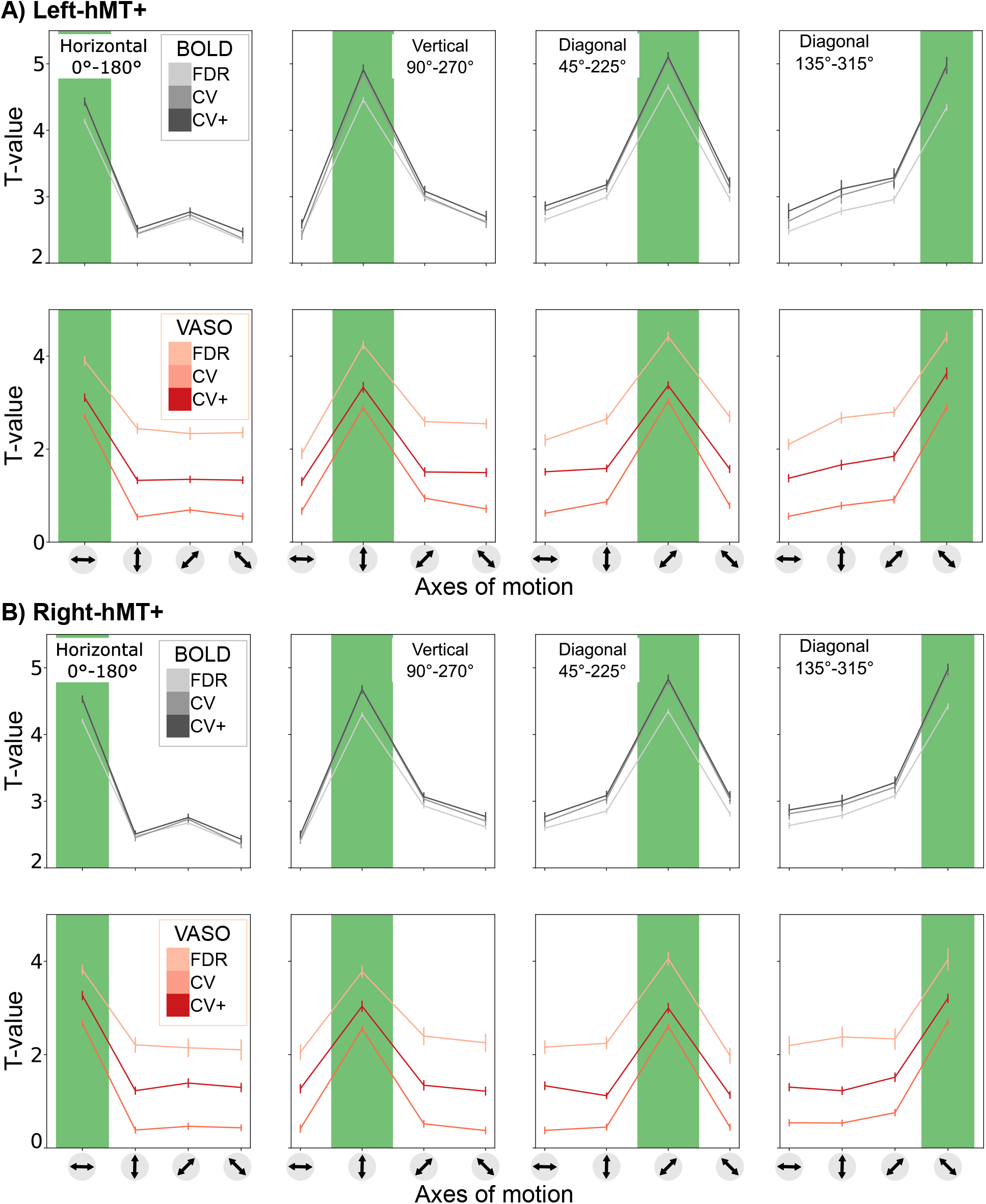
Group average axis-of-motion selectivity tuning curves for each axis-of-motion for both BOLD and VASO contrast in the (A) left hMT+ and (B) right hMT+. Tuning curves are evaluated for FDR, CV and CV+ voxels. The plots depict the mean and the standard error of all voxels for each category.

For the BOLD contrast, varying the voxel selection method results in a subtle progressive increase in tuning selectivity index from the FDR to the CV and CV+ voxels (see **Table 2**). **Figure 3** shows that the t-value of the preferred condition increases, while t-values for the non-preferred conditions decrease. We don’t observe substantial differences when comparing the CV with CV+ BOLD voxels, since the voxel’s reduction rate was always less than 10%. Our BOLD results consistently show the tuning selectivity of hMT+ being invariant to the voxel selection procedure.

**Table 2:**
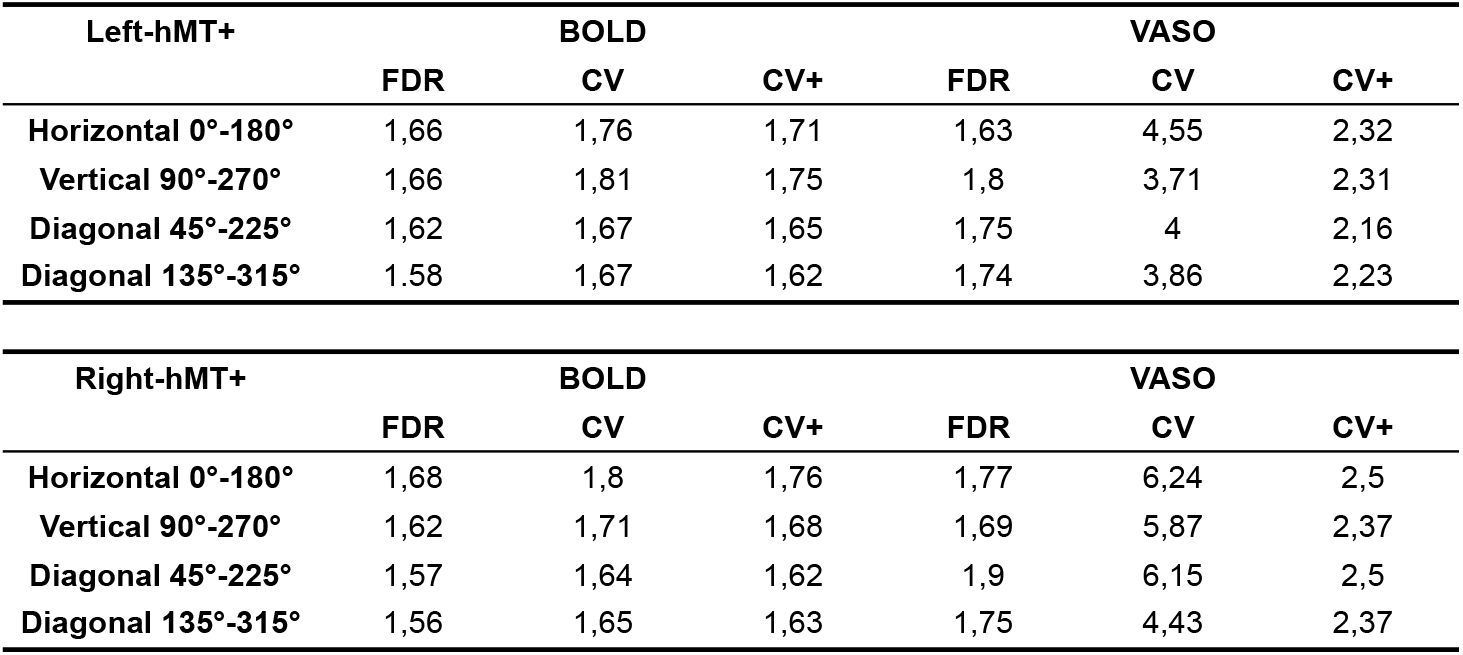
Tuning Selectivity Index reported for both BOLD and VASO and each voxel selection method (FDR, CV, CV+).

| Left-hMT+ | BOLD |  |  | VASO |  |  |
| --- | --- | --- | --- | --- | --- | --- |
|  | FDR | CV | CV+ | FDR | CV | CV+ |
| Horizontal 0°-180° | 1,66 | 1,76 | 1,71 | 1,63 | 4,55 | 2,32 |
| Vertical 90°-270° | 1,66 | 1,81 | 1,75 | 1,8 | 3,71 | 2,31 |
| Diagonal 45°-225° | 1,62 | 1,67 | 1,65 | 1,75 | 4 | 2,16 |
| Diagonal 135°-315° | 1,58 | 1,67 | 1,62 | 1,74 | 3,86 | 2,23 |

| Right-hMT+ | BOLD |  |  | VASO |  |  |
| --- | --- | --- | --- | --- | --- | --- |
|  | FDR | CV | CV+ | FDR | CV | CV+ |
| Horizontal 0°-180° | 1,68 | 1,8 | 1,76 | 1,77 | 6,24 | 2,5 |
| Vertical 90°-270° | 1,62 | 1,71 | 1,68 | 1,69 | 5,87 | 2,37 |
| Diagonal 45°-225° | 1,57 | 1,64 | 1,62 | 1,9 | 6,15 | 2,5 |
| Diagonal 135°-315° | 1,56 | 1,65 | 1,63 | 1,75 | 4,43 | 2,37 |

For the VASO contrast, varying the voxel selection method from FDR to CV+ does not result in a progressive increase in tuning selectivity index. The tuning curves from the FDR voxels are characterized by the highest t-value range in both preferred and not preferred conditions. As a consequence, the tuning selectivity index is the smallest among the three sets and comparable with the ones computed for the BOLD contrast. The statistical threshold q(FDR)<0.05 might be too conservative for the low sensitivity of the VASO contrast and only captures a subset of tuned voxels that are similarly activated in both contrasts. The tuning curves from the CV voxels are characterized by the highest tuning selectivity index. The percentage of surviving CV voxels around 24.8% (left-hMT+) and 19.2% (right-hMT+) was comparable with 35.8% (left-hMT+) and 37.6% (right-hMT+) for the BOLD contrast, suggesting that the partial recovery of the low sensitivity for the VASO contrast comes together with an increased tuning selectivity (see **Table 2**). However, when considering the VASO tuning curves using CV+ voxels, the tuning selectivity index is still higher than VASO FDR voxels, but it is lower when compared to VASO CV voxels. This effect might be due to the presence of negative t-values in the CV voxels, which might pull towards zero the t-values of the non-dominant conditions, inflating the tuning selectivity index. Although we found differences in tuning selectivity depending on the voxel selection, our results show tuning selectivity for both BOLD and VASO tuning curves with a slightly stronger selectivity effect for the VASO contrast.

### 3.2. Voxel-wise sensitivity and specificity metrics quantify differences in imaging contrast

We computed voxel-wise sensitivity and specificity for all FDR, CV, CV+ tuned voxels for both BOLD and VASO contrast in both hemispheres. As observed in **Figure 4** characteristic sensitivity-specificity scatter plots were found for both BOLD and VASO contrast. For all participants, we observed that BOLD cross-validated voxels are overall more sensitive (spanning a wider sensitivity range) and less specific (spanning a narrower specificity range) compared to VASO voxels. In line with our tuning curve results, the choice of the voxel selection does not affect the shape of both BOLD and VASO scatterplots. However, the strongest BOLD and VASO difference in sensitivity and specificity ranges are mostly observed for CV and CV+ voxels. Again, the FDR VASO voxels seem to be a subset of BOLD voxels.

**Figure 4:**
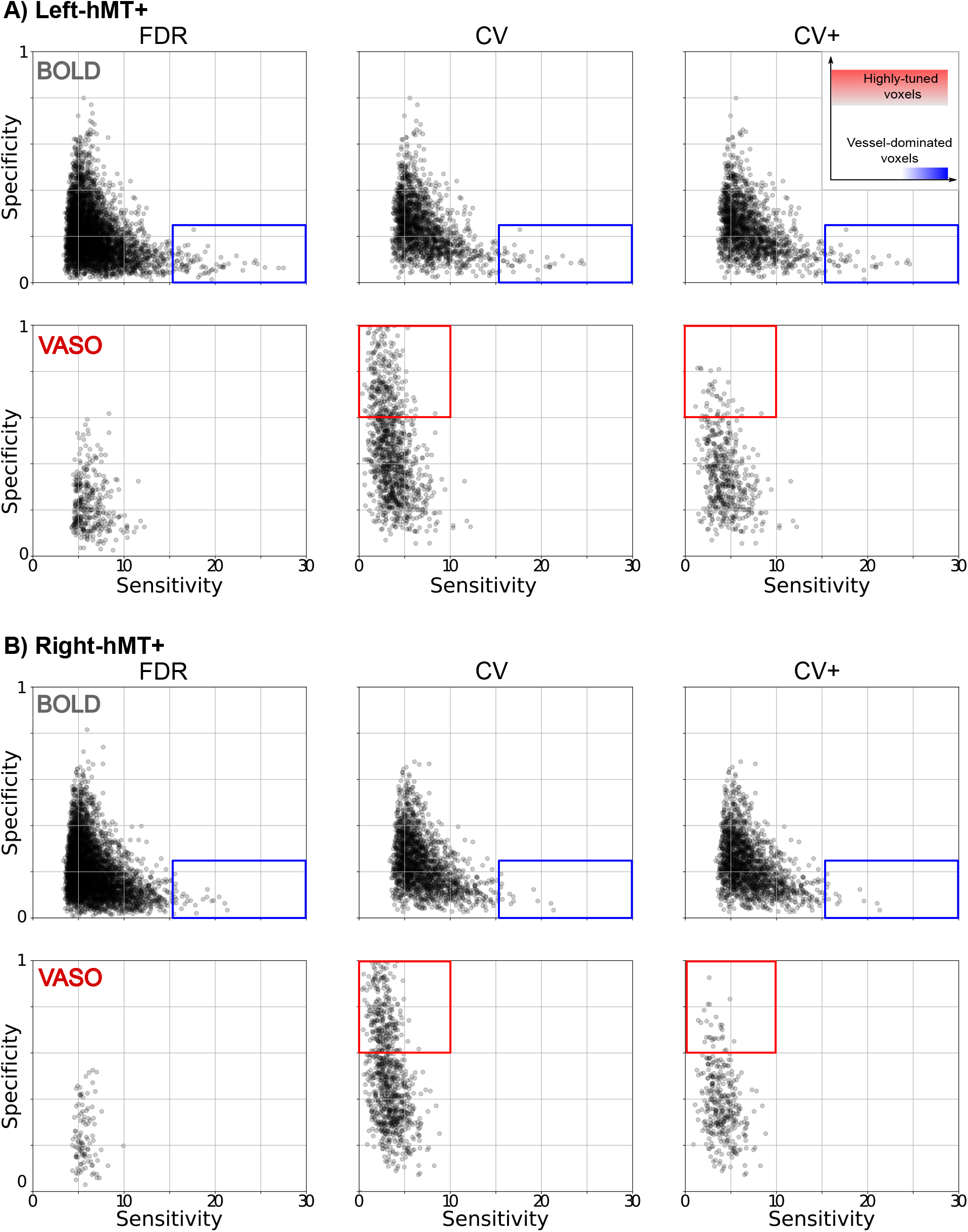
Group result for sensitivity and specificity scatterplot for both BOLD and VASO contrast in (A) left hMT+ and (B) right hMT+ for. FDR (right column), CV (middle column) and CV+ (right column) voxel sets. The inset at the upper right corner shows in a qualitative way how different regions of the space are associated with different voxel characteristics. Rectangles highlight main differences between contrast types and voxel selection strategy. The red rectangle encompasses highly tuned voxels, whereas the blue area shows vessel-dominated voxels.

Our results are in agreement with recent findings (Beckett et al., 2020; Huber et al., 2017) (see also **Supplementary Figure 2 and 3**) and confirm that, despite its reduced signal sensitivity, the VASO contrast provides greater specificity to the neuronal origin of the fMRI signal. Interestingly, our new local sensitivity and specificity measures offer a voxel-wise perspective on the draining veins effect: the vessel-dominated voxels responsible for this effect can be straightforwardly detected and localized. As shown for an exemplary participant in **Figure 5**, the combination of high sensitivity and low specificity help detecting vessel-dominated voxels (highlighted by green arrows) similarly to decreased BOLD T_2_*-weighted EPI intensity that has been shown to be a robust marker of vascular effects (Kay et al., 2019). A lower number of vessel-dominated voxels are found for VASO contrast compared to BOLD, demonstrating the reduced signal contamination by draining veins and its improved specificity. In **Figure 6**, we project the vessels-dominated voxels shown in **Figure 5** in the flat laminar-resolved domain. This representation provides a more convenient visual representation to evaluate voxel localization across cortical depths. As shown in **Figure 6**, the vessel-dominated voxels appear mostly in the superficial layers and disappear in the deep layers. This result demonstrates BOLD signal contamination by pial vasculature that lies above the cortex. The same phenomenon is strongly attenuated in VASO signal.

**Figure 5:**
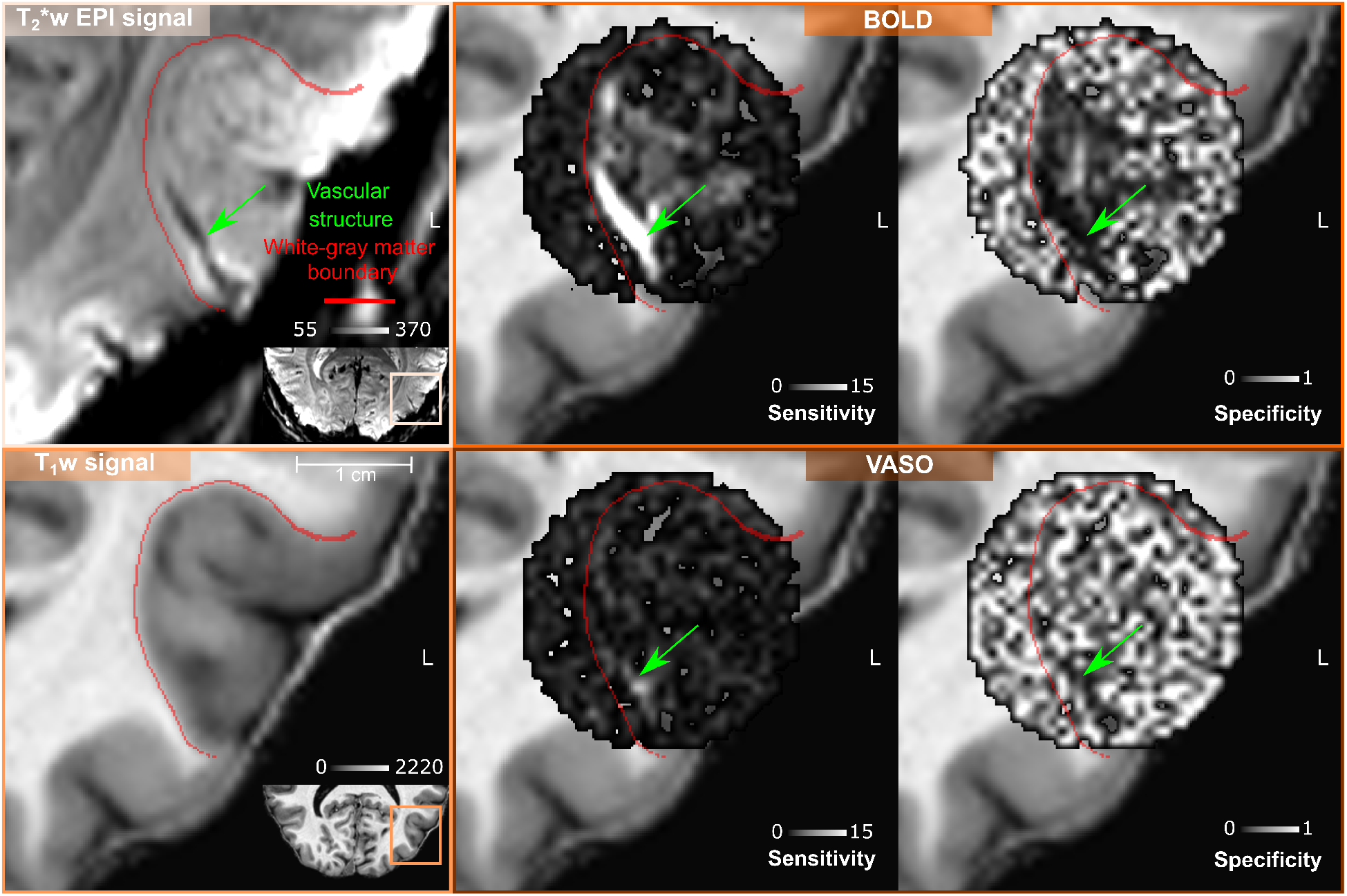
Volume-based visualization of voxel-wise sensitivity and specificity maps for one example participant (sub-01) in the left hMT+ ROI. The same axial slice is shown for both T_2_*-w EPI signal and T_1_-w signal (nominal resolution 0.2 iso mm). The green arrow points to a vascular structure detectable only through T_2_*-w EPI intensity as a dark spot (reflecting static susceptibility effects). High sensitivity and low specificity characterizes all the voxels belonging to this vascular structure (green arrows) for BOLD contrast. The detectability of this vascular structure is less evident for VASO contrast, as expected.

**Figure 6:**
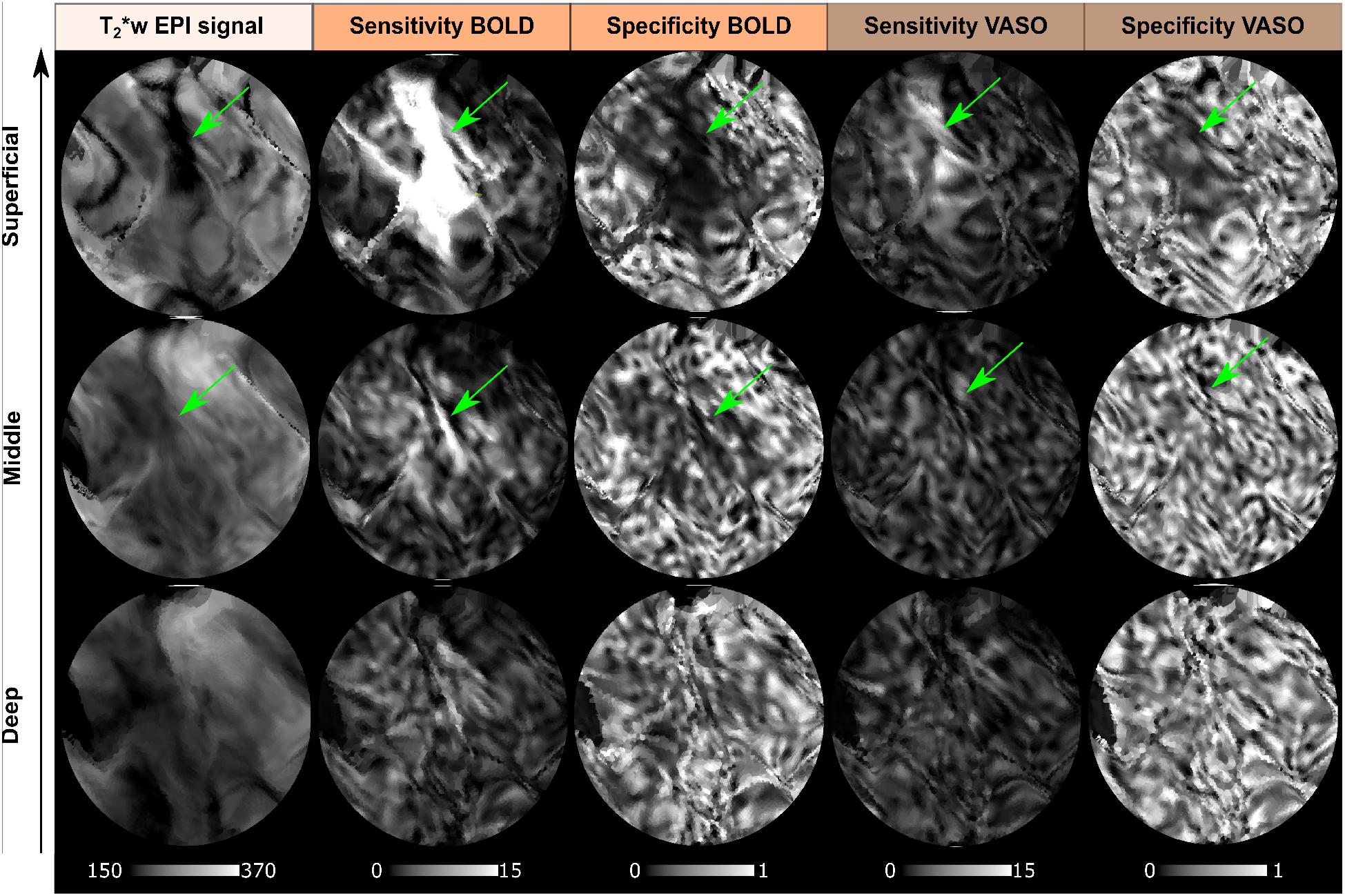
Depth-dependent T_2_*-w EPI intensity, voxel-wise sensitivity and specificity maps for BOLD and VASO contrast showed for the flattened left hMT+ ROI for one example participant (sub-01) (nominal resolution 0.05 mm iso.). Note that in this figure we project the same vessels-dominated voxels shown in **Figure 5** in the flat laminar-resolved domain. The vascular structure highlighted by the green arrow corresponding to the dark spot in T_2_*-w EPI intensity is clearly visible at superficial layers but not at middle or deep layers, which indicates that it is a pial vein. High sensitivity and low specificity are in agreement with the spatial displacement of darkness of T_2_*-w EPI intensity.

### 3.3. Quantification of the columnar organization of axes of motion tuned voxels

We explored the spatial organization of axis-of-motion tuned voxels for both BOLD and VASO contrast by using our new searchlight algorithm for column detection. We evaluated both BOLD and VASO columnarity maps as a function of the columnarity index. We used the flattening algorithm (Gulban et al., 2022) and 3D rendering visualization tools (Sullivan and Kaszynski, 2019) to show our columnar results. The spatial organization of the columns can be fully observed in the rotated animations accompanying the **Figure 7**: https://doi.org/10.6084/m9.figshare.20393667.

**Figure 7:**
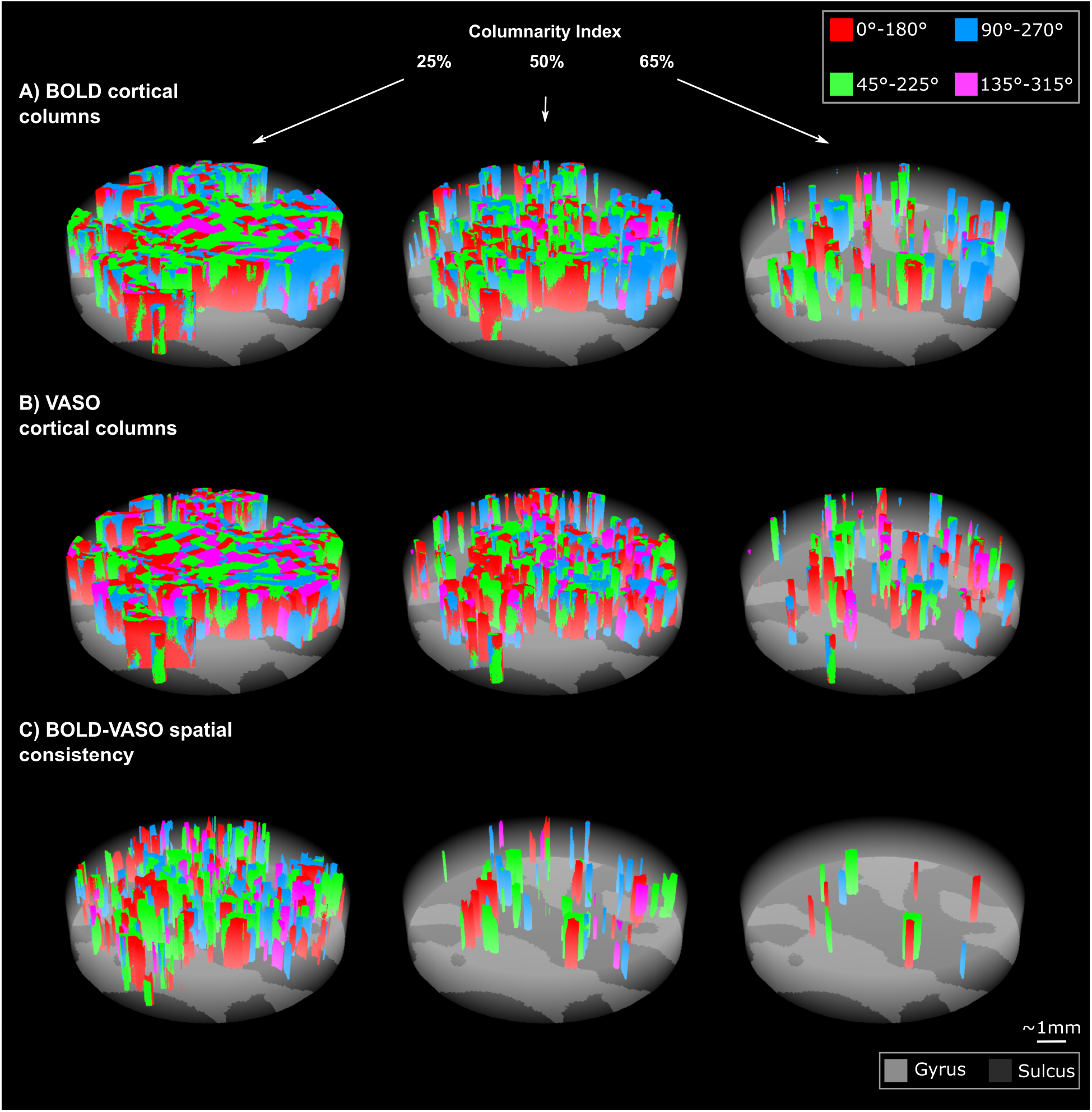
Examples of BOLD (A) and VASO (B) axes-of-motion cortical columns for one example participant (sub-01) at 25%, 50%, 65% of the columnarity index. C) BOLD-VASO spatial consistency plots show only axes of motion columns that are detected in both BOLD and VASO contrasts for each columnarity threshold (see animations here: https://doi.org/10.6084/m9.figshare.20393667.

In **Figure 7A, B** we show BOLD and VASO axis-of-motion columnar results for one representative participant (left-hMT+) for three values of the columnarity map 25%, 50%, 65%. Qualitatively, we observe that hMT+ is mostly organized by ‘small’ columns or ‘patch of columns’ occupying a quarter or a half of the whole cortical depth. For these columnarity index thresholds, we find that only a part of the columnar organization is preserved in both BOLD and VASO contrast (**Figure 7C**). The density of the axis-of-motion columns decreases with a linear increase of the columnarity index for both BOLD and VASO.

To derive a quantitative description of the columnar organization of hMT+ with respect to the axis-of-motion stimuli, we summarized the group results of hMT+ in **Figure 8**. According to our local measures of sensitivity and specificity, we confirm that VASO columnarity results are always more specific and less sensitive than BOLD columnarity results regardless of the choice of the columnarity index (**Figure 8A,B (i-ii)**). Interestingly, we found that for a columnarity index of 40% the percentage of columnar volume reaches a peak for both BOLD and VASO contrast (**Figure 8A,B (iii)**). At the same time, more than 50% of the columnarity maps are consistently found in both BOLD and VASO columnar results (**Figure 8A,B (iv)**). Our results suggest that the selectivity of hMT+ is mostly organized in ‘patches of columns’, and only partially organized in geometrically-defined columns spanning the whole cortical depth.

**Figure 8:**
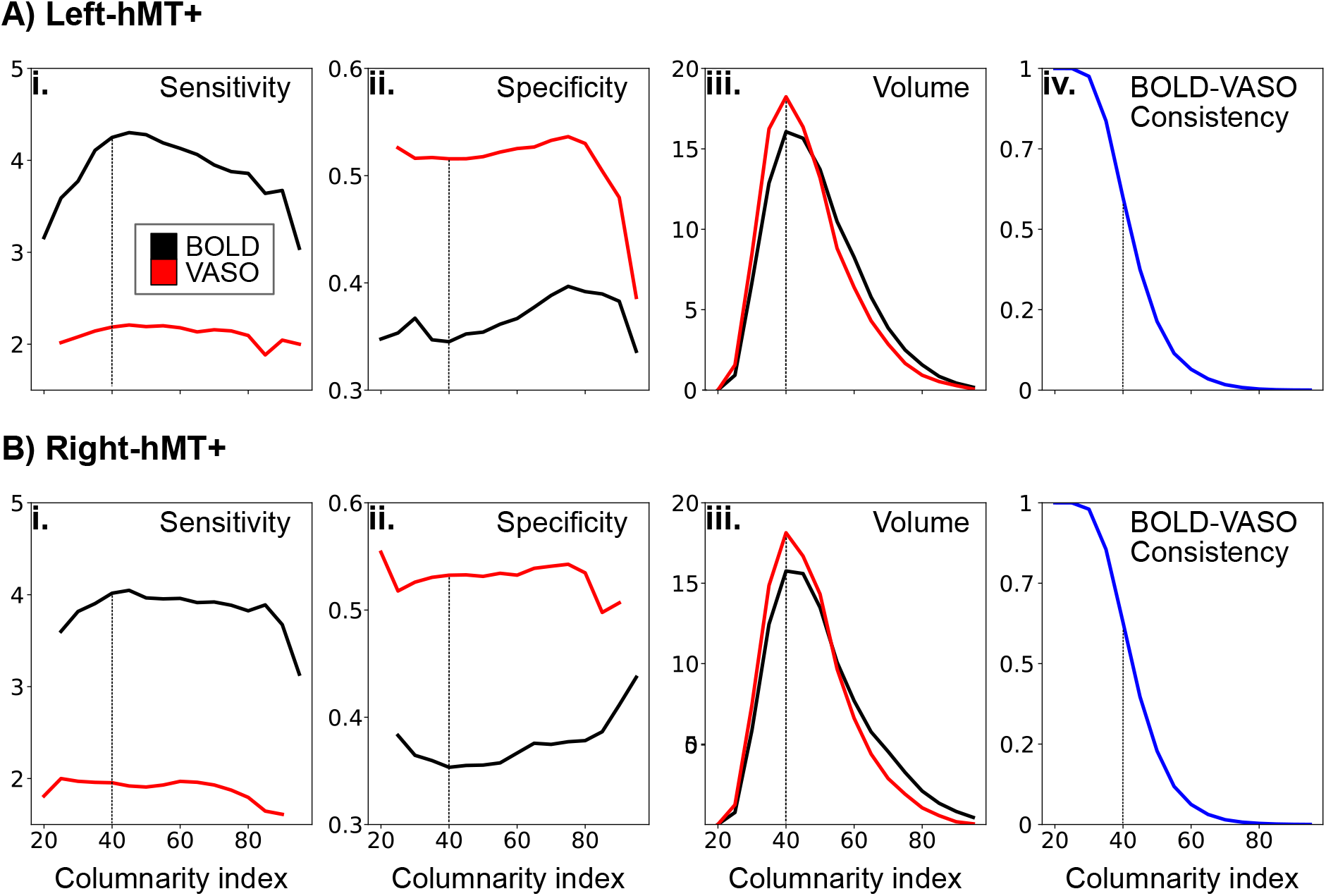
Descriptive statistics for both BOLD and VASO columnarity maps in (A) left hMT+ and (B) right hMT+. Average of voxel-wise sensitivity (i) and specificity (ii) and percentage of columnar hMT+ volume (iii) is evaluated for both BOLD (black line) and VASO (red line) as a function of the columnarity index. (iv) Average of BOLD-VASO consistency quantified for each value of the columnarity index. A vertical dotted line is drawn for a columnarity index of 40, corresponding to the peak of columnar volume (iii).

Finally, we compare our empirical results with the columnar results derived from the ‘random’ and ‘ideal columnar’ benchmark results and summarize them in **Figure 9**. In **Supplementary Figure 4** we compare columnarity index’s probability density function (PDF) of empirical and benchmark data for each participant. As we can see for the group results in **Figure 9**, the median of the distribution of the columnarity index for both BOLD and VASO is always in between the median of the distribution of the columnarity index of the ‘random’ and the ‘ideal’ benchmark datasets.

**Figure 9:**
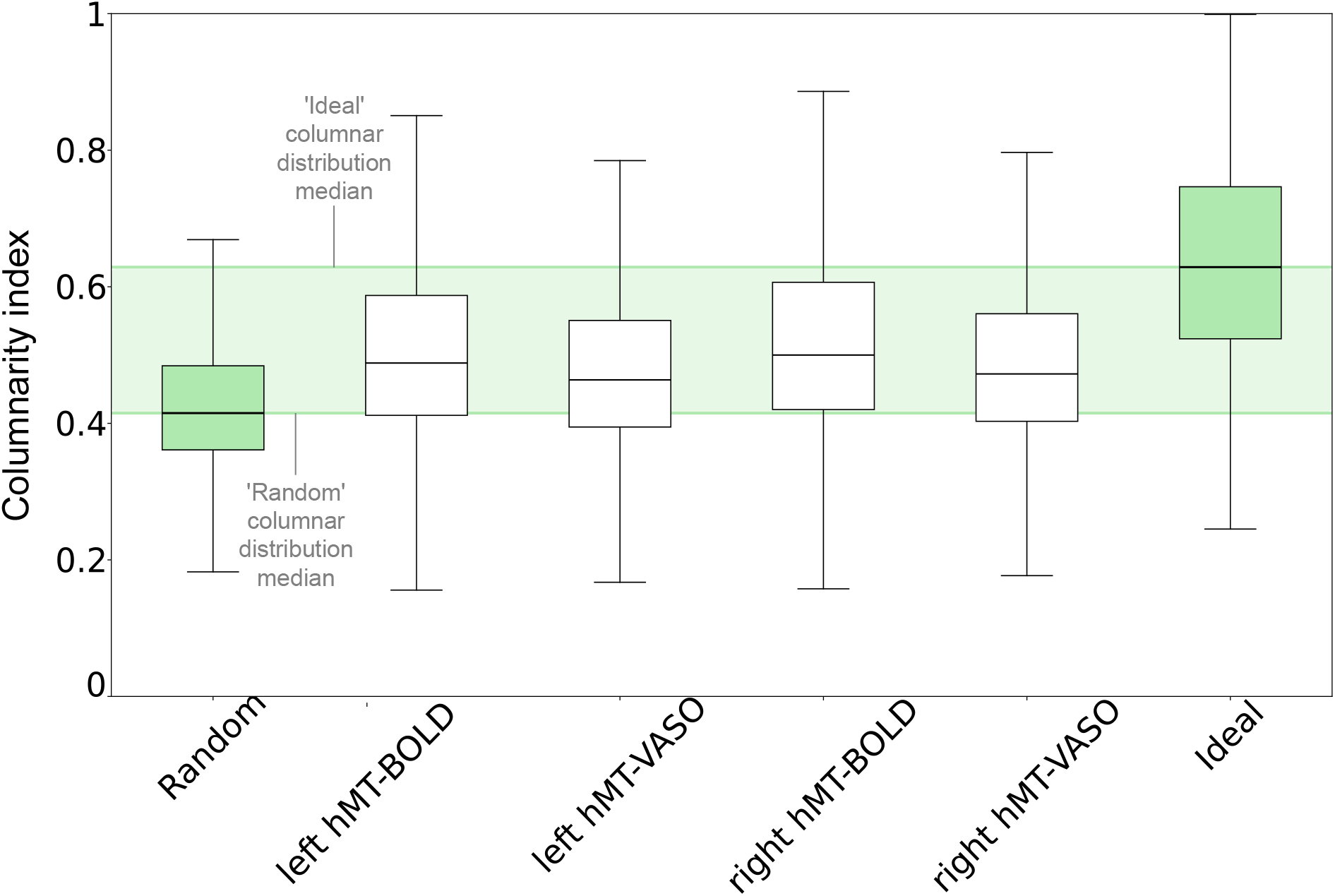
Comparing columnarity index distributions between empirical and benchmark datasets. Empirical BOLD and VASO group distributions of left-hMT+ and right-hMT+ are reported as white boxplots. ‘Random’ and ‘ideal’ columnarity index distributions are shown in green boxplots on the left and right side, respectively. Boxplots extend from the first quartile to the third quartile, with a line at the median. The whiskers extend from the box by 1.5x the iter-quartile range. Green shaded bands encompass the upper and lower distribution of the random and ideal benchmark sets. See **Supplementary Figure 4** for a detailed version with individual participant results.

## 4. Discussion

### 4.1. Overview of the results

This study shows for the first time the feasibility of mapping axes of motion cortical columns in the extrastriate area hMT+ in living humans using CBV-VASO fMRI at 7T. The SS-SI VASO sequence with 3D EPI readout enabled us to simultaneously acquire BOLD and VASO responses in hMT+. Therefore, we separately applied the same analysis pipeline to both VASO and BOLD fMRI data (Huber et al., 2015, 2017; Oliveira et al., 2022; Oliveira et al.). In this regard, the development of new local metrics of sensitivity and specificity was pivotal to quantitatively compare and interpret results found with VASO and BOLD contrast. Our metrics not only confirmed the higher specificity and lower sensitivity of VASO compared to BOLD voxels (Beckett et al., 2020), but also allowed us to clearly highlight pial vein effects in BOLD contrast. Finally, the new searchlight algorithm for functional column detection provides a unique framework to investigate mesoscopic cortical features with its improved quantifiability and comparability with respect to previously used methodologies (Yacoub et al., 2008; Zimmermann et al., 2011; De Martino et al., 2013, 2015; Schneider et al., 2019). The columnarity map computed by our algorithm provides a full representation of the 3D functional organization of hMT+ allowing a transparent evaluation of columnarity.

#### 4.1.1. *Local* vs *global sensitivity and specificity metrics*

Conventionally, global measures of sensitivity and specificity (Huber et al., 2017; Beckett et al., 2020) are used to compare different contrast mechanisms in terms of draining veins effects. As described in (Beckett et al., 2020) those global indices are computed by fitting a linear regression model to layer profiles (see **Supplementary Figure 3**). In this work, we extended the concept of sensitivity and specificity from a global to a voxel-wise scale, by exploiting the tuning property of hMT+ voxels. By visualizing our sensitivity and specificity maps, we observed that untuned voxels show very low specificity. Voxels close to pial vessels show a very low specificity and very high sensitivity in combination with a low T_2_*-w EPI signal as an additional independent diagnostic marker. For the data of the participant reported in **Figure 5** and in **Figure 6**, we clearly showed BOLD voxels sampling a pial vessel. The VASO contrast is expected to be not sensitive to macrovasculature. However, we still observe a smaller number of voxels with the aforementioned properties. Two possible candidate mechanisms can be leading to this phenomenon: a flow-dependent vein effect as described in (Huber, 2015) or a dilatation effect of big arteries on the pial surface (Kim et al., 2013), which is still disputed. We believe that our new local metrics of sensitivity and specificity provide an alternative way to evaluate the draining veins effect and to visualize pial vessels responsible for it.

#### 4.1.2. General perspective on cortical columns

While the existence of direction selective neurons within area MT is a well-established feature in primates (Albright, 1984) and cats (Hubel and Wiesel, 1962, 1965) only preliminary fMRI evidence has been reported in humans (Zimmermann et al., 2011; De Martino et al., 2013). Our findings provide strong evidence for a functional columnar organization of axis-of-motion features also in human MT+. The obtained columnarity maps, however, indicate that functional clusters are only partially in line with proposed ideal (anatomical) column models (Mountcastle, 1956) that are assumed to penetrate vertically from the pial surface to the white matter boundary in a regular manner. The hypothesis of 3D columns was mostly investigated by multi-unit electrophysiological recordings in animals. Pioneering results from Hubel and Wiesel (1962, 1965); Albright et al. (1984), were put in perspective by reports showing that the preferred orientations were not necessarily represented in a columnar fashion in animal visual cortex (Bauer et al., 1983, 1989; Berman et al., 1987). In particular, Bauer et al. (1983) reported that the preferred orientations jump by 90° along the vertical track of the electrode penetration in cat area 18. It’s worth noting that this technique comes with the cumbersome task of accurately tracing the penetrating electrodes together with the limited sampling space, affecting the robustness of those results (Horton and Adams, 2005). Later findings (Tanaka et al., 2011) provided a new perspective on the conventional columnar view of orientation representation of the visual cortex: simulations based on a 3D self-organized model predicted the occurrence of direction reversal in columns along the cortical depth dimension being proportional to the curvature of the cortex, and orientation columns having wedge-like shape when sampling gyri or sulci. The same study confirmed these theoretical predictions with multi-slice, high-resolution functional MRI in cat areas 17 and 18. These mechanisms were also discussed by (Zimmermann et al., 2011) to explain the variability of iso-oriented feature maps across cortical depths observed with high-resolution fMRI in humans. Recently, Nakamichi et al. (2018) proposed a new explanation for the variability across cortical depths based on functional optical coherence tomography in cats: their results show that the 3D structure of orientation columns were heavily distorted around pinwheel centers. Orientation singularities were rarely straight solid bars connecting pinwheel centers, instead they typically ran inside the cortex creating “singularity strings” with peculiar trajectories. Besides findings challenging a simplistic view of columnar organization (Rakic, 2008), the functional relevance of cortical columns has also been debated (Horton and Adams, 2005; Haueis, 2021). Whether cortical columns “are a structure without a function” (Horton and Adams, 2005) or not, the presence of functional clusters with groups of neurons that share similar tuning properties allows high-resolution human fMRI studies to reveal coding principles of the brain, which would otherwise only be possible at microscopic resolution.

Our columnar results are to our knowledge the first to provide a quantitative columnarity map to characterize the functional organization of axis-of-motion features in hMT+ exploiting the benefits of CBV-based fMRI at 7T. Our new algorithm for column detection not only improved the quantifiability and comparability with other studies, but it also takes into account the curvature effect (Tanaka et al., 2011). However, we found that the spatial consistency between VASO and BOLD columnar results decreased as a function of the columnarity index. We believe that this variability is due to an interplay of two effects. On the one hand, the differences in terms of sensitivity but especially specificity between VASO and BOLD would mostly affect the columnar results when a conservative columnarity index threshold is applied. On the other hand, the aforementioned theory of variability of orientation selectivity across cortical depths could explain the trend of spatial consistency.

Future CBV-sensitive fMRI studies with higher spatial resolution (e.g. < 0.5 mm isotropic) and improved sensitivity would be able to investigate the functional organization of hMT+ in more detail that would be likely sufficient to resolve direction-of-motion columns instead of the larger axis-of-motion columns. Such higher-resolution fMRI studies might also reveal direct evidence for pinwheels and their effect on columnarity. The improved quantifiability provided by our new method for column detection will make the comparison of our results with future studies straightforward. Furthermore, our approach and methodological developments are generalizable and applicable to other human brain areas where similar mesoscopic research questions are addressed.

## Data and code availability statement

Analysis codes are available on github: https://github.com/27-apizzuti/AOM-VASO_project. A full overview of our processing pipeline is available here: https://github.com/27-apizzuti/AOM-VASO_project/tree/main/diagrams. Data will be made available upon manuscript acceptance.

## Author Contributions

According to the CRediT system (https://casrai.org/credit/)

**Conceptualization:** A.P., R.H., O.F.G., R.G.

**Methodology:** A.P., O.F.G., J.P., R.G.

**Software:** A.P., O.F.G.

**Validation:** A.P., R.H., O.F.G., R.G.

**Formal Analysis:** A.P.

**Investigation:** A.P., R.H., A.B.A.

**Resources:** A.P., R.H., O.F.G., A.B.A., J.P., R.G.

**Data curation:** A.P., A.B.A.

**Writing – original draft:** A.P.

**Writing – review & editing:** A.P., R.H., O.F.G., A.B.A., J.P., R.G.

**Visualization:** A.P., R.H., O.F.G., R.G.

**Supervision:** R.H., O.F.G., J.P., R.G.

**Project administration:** R.H., J.P., R.G.

**Funding acquisition:** R.G.

## Acknowledgements

This project was funded by the EU-project H2020-860563 euSNN and the European Union’s Horizon 2020 Framework Programme for Research and Innovation under the Specific Grant Agreement No. 945539 (Human Brain Project SGA3). Laurentius Huber was funded from the NWO VENI project 016.Veni.198.032. The sequence used here is based on code kindly written and provided by Benedikt A. Poser. Data was acquired at Scannexus (Maastricht, the Netherlands). We thank Chris Wiggins for providing the 3rd order shimming tools. We thank David Feinburg for valuable discussions on the columnar results. We would also like to thank *Maastricht Layer-Seminar* members Sebastian Dresbach, Lonike Faes, Miriam Heynckes, Kenshu Koiso, and Yawen Wang for discussions throughout the project.

## Supplementary Material

**Supplementary Figure 1:**
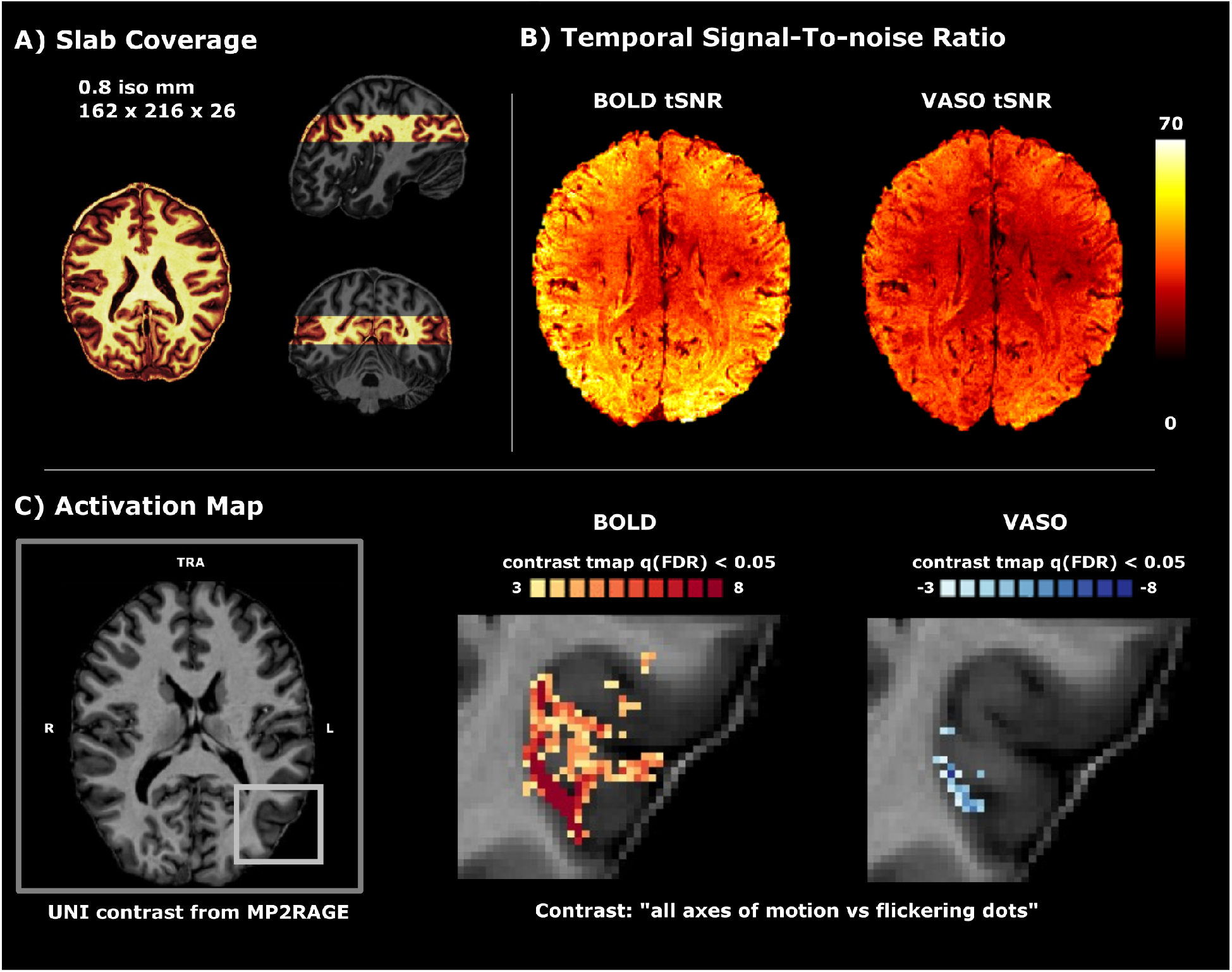
SS-SI VASO functional images for an example participant (sub-01). A) Slab coverage (in warm colors) relative to a whole brain MP2RAGE. B) Temporal signal-to-noise ratio. C) Activation map for BOLD and VASO time series for the contrast: “all axes of motion vs flickering dots” (q(False Discovery Rate) < 0.05). for the left-hMT+.

**Supplementary Figure 2:**
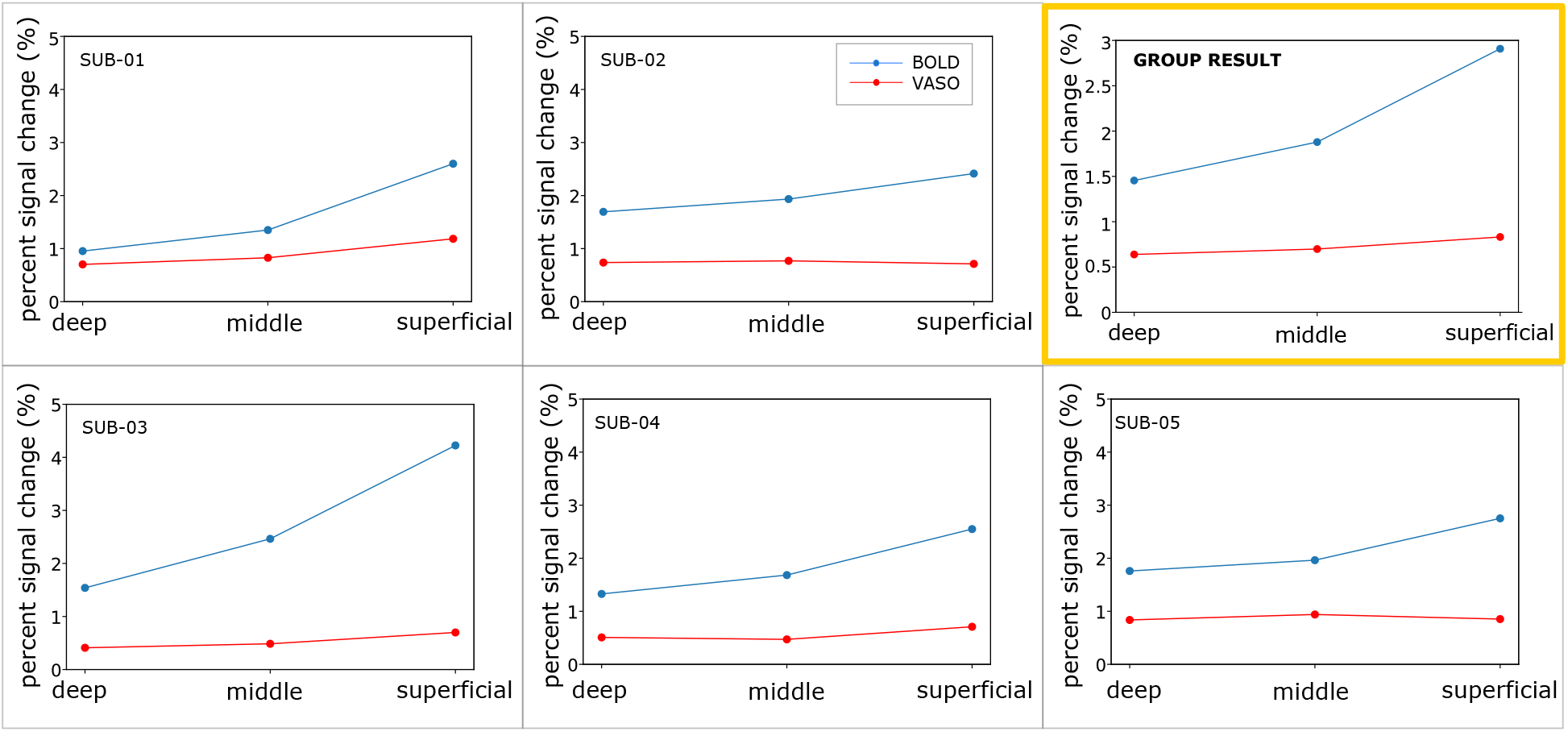
Layer profiles across three cortical depths: deep (25%), middle (50%), superficial (75%) with respect to the white-gray matter boundary in the left hMT+. For each cortical depth, the averaged percent signal change of all cross-validated voxels is reported for both BOLD and VASO. The yellow box shows the group result of all five participants. See **Supplementary Figure 3A** for a detailed version of this figure showing individual participants and conditions.

**Supplementary Figure 3:**
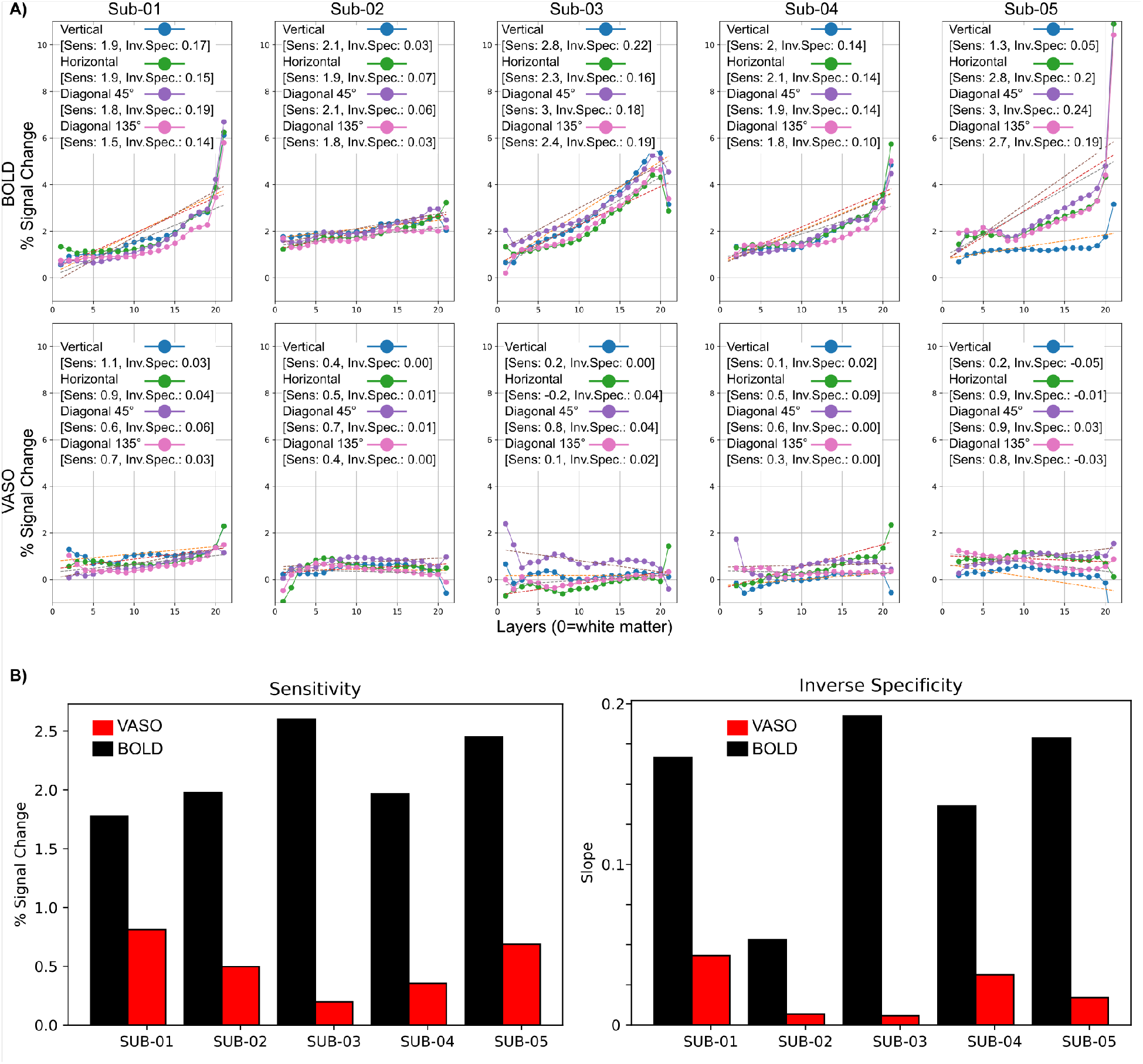
Global metrics of sensitivity and inverse specificity computed for each subject (columns), for both BOLD (first row) and VASO contrast (second row) considering cross-validated voxels in the left hMT+. Each subplot shows single layer profiles for each axis-of-motion condition. The square brackets report global sensitivity and inverse measure of global specificity (Beckett et al., 2020). Condensed version of this panel showing average stimulation conditions can be seen in **Supplementary Figure 2**. B) Global metrics of sensitivity and inverse specificity (Beckett et al., 2020) computed for each participant, for both BOLD (black bar) and VASO (red bar). Each barplot depicts the mean value evaluated across the four axes of motion.

**Supplementary Figure 4:**
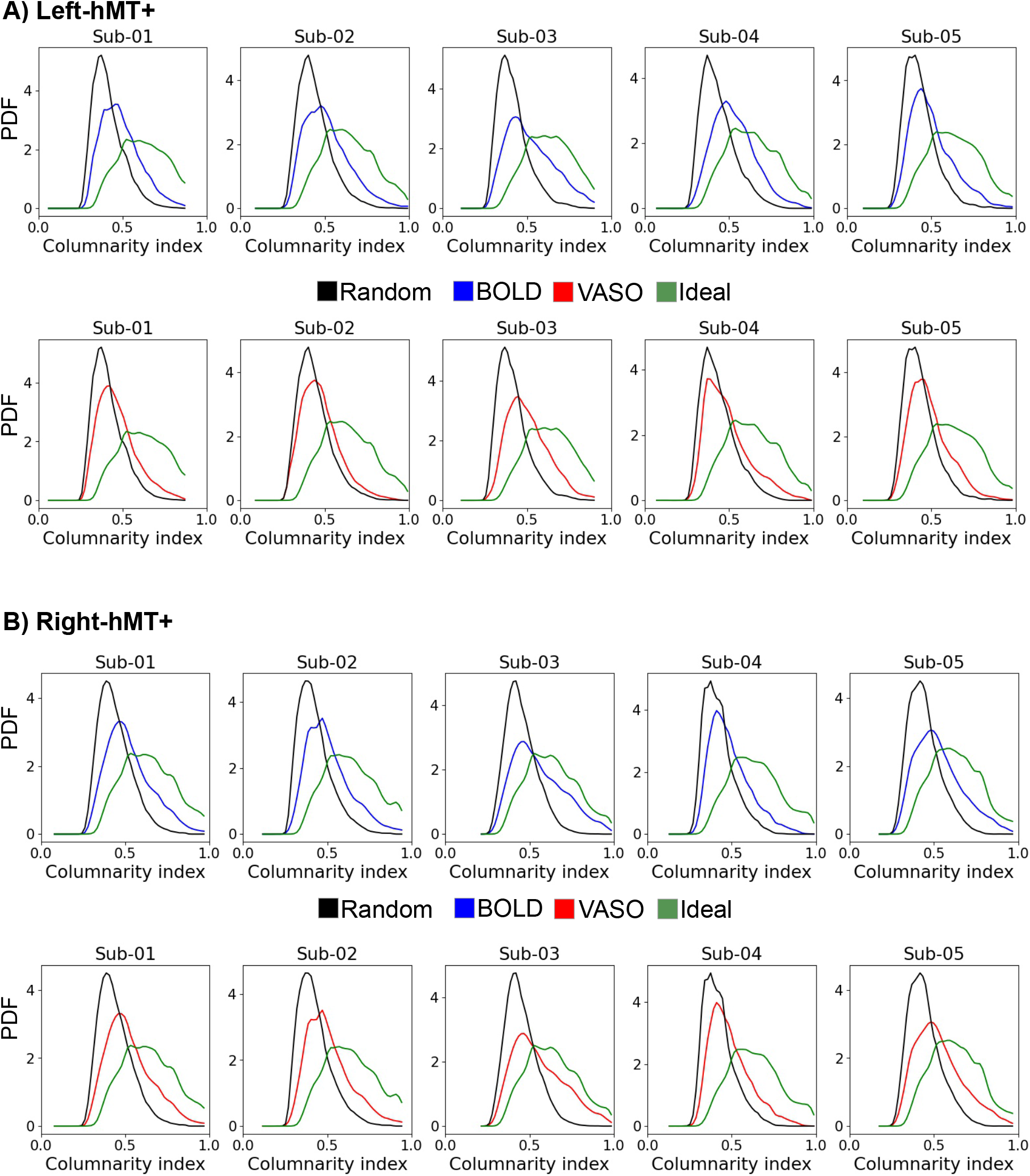
Comparing columnarity’s index probability density function (PDF) of empirical and benchmark data for each participant. First rows of panel A and B show BOLD (blue color), ‘random’ and ‘ideal’ PDFs. Second rows of panel A and B show VASO (red color), ‘random’ and ‘ideal’ PDFs. Condensed version of this figure showing group results can be seen in **Figure 9**.

